# Hematopoietic differentiation is characterized by a transient peak of entropy at a single cell level

**DOI:** 10.1101/2021.04.30.442092

**Authors:** Charles Dussiau, Agathe Boussaroque, Mathilde Gaillard, Clotilde Bravetti, Laila Zaroili, Camille Knosp, Chloé Friedrich, Philippe Asquier, Lise Willems, Laurent Quint, Didier Bouscary, Michaela Fontenay, Thibault Espinasse, Adriana Plesa, Pierre Sujobert, Olivier Gandrillon, Olivier Kosmider

**Author notes:** These authors contributed equally to this work (Pierre Sujobert, Olivier Gandrillon, Olivier Kosmider). Corresponding author: Mail (OK).

## Abstract

Hematopoietic differentiation has been metaphorically represented as linear trajectories with discrete steps from hematopoietic stem cells to mature cells. While the transcriptional state of cells at the beginning or at the end of these trajectories are well described from bulk analysis, what happens in the intermediate states has remained elusive until the use of single cell approaches. Applying Shannon entropy to measure cell-to-cell variability among cells at the same stage of differentiation, we observed a transient peak of gene expression variability in all the hematopoietic differentiation pathways. Strikingly, genes with the highest entropy variation in a given differentiation pathway matched genes known as pathway-specific, whereas genes with the highest expression variation were common to all pathways. Finally, we showed that the level of cell-to-cell variation is increased in the most immature compartment of hematopoiesis in myelodysplastic syndromes. These data suggest that differentiation could be better conceptualized as a dynamical stochastic process with a transient stage of cellular indetermination.

## Introduction

Complex biological processes such as development or differentiation are often conceptualized as the execution of a program encoded in the genome. However, the existence of random process is increasingly recognized, especially at the level of gene expression, a fundamentally stochastic process, leading to cell-to-cell variations in mRNA and protein levels (*1*).

Hematopoiesis is a finely regulated process by which hematopoietic stem cells (HSCs) give rise to mature blood cells that belong to myeloid or lymphoid lineages. HSC differentiation toward a lineage is thought of as a continuous process through a series of increasingly committed progenitors (*2*) along one of several trajectories leading to the production of the various mature blood cells. Whether the process of hematopoietic stem cell commitment is instructive or stochastic has long been the subject of controversies (*3, 4*). According to the instructive model, HSC receive external signals such as cytokines which actively induces them to differentiate into a given lineage(*5–7*). On the contrary, the stochastic model proposes that spontaneous stochastic variations of cell phenotype are followed by a selection step driven by cytokines (*8, 9*). Following the instructive model, cell-to-cell phenotypic variability should be limited and should not vary during differentiation, since all of the cells committed to a given lineage are following the same instructions. On the contrary, the stochastic model predicts that cell-to-cell variability should increase to reach a maximum at the stage where the selection occurs.

Until recently, the analysis of hematopoietic differentiation has relied on transcriptomic analyses of bulk populations of sorted cells defined by their common phenotype, or on the analysis of only a few markers captured at the single cell level by flow cytometry. With the technological breakthrough of single cell transcriptomics (*10*), we are now able to study the whole transcriptome of each cell during the differentiation process, hence allowing a comprehensive measure of cell-to-cell variability. Based on the quantification of a limited number of mRNA, we have shown that a surge in gene expression variability occurs during avian erythropoiesis in an *in vitro* model (*11*). This was further confirmed in experimental differentiation models *in vitro* (*12–15*) or in animals (*16, 17*). Moreover, variability in gene expression has been suggested to play a causal role in cell differentiation leading many studies to use single cells approaches(*15, 18*). However, these observations were limited either by the number of genes analyzed (*11, 12, 16, 17*), or by the question of the physiological relevance of the established cell lines used (*14*).

Here we used the conceptual framework of Shannon entropy as a proxy of cell-to cell variation, to analyze for each gene cell-to-cell gene expression variability during HSC differentiation in normal scRNA-seq human bone marrow datasets. We observed a peak of entropy along all the differentiation pathways (erythroid, granulocytic, dendritic and B lymphoid). Of note, genes with the highest entropy variation in a given differentiation pathway corresponded to genes known as pathway-specific, whereas genes with the highest expression variation were common to all pathways. Finally, applying this approach to analyze bone marrow from patients with myelodysplastic syndromes, which are characterized by ineffective differentiation, we observed a higher level of entropy in the most immature states in the affected patients.

## Results

### A peak in cell-to-cell gene expression variability is a common feature of all hematopoietic differentiation lineages

In order to study stochastic gene expression during normal hematopoiesis, we analyzed public scRNA-Seq data of 15962 genes in 12602 mononuclear cells derived from the bone marrow of a healthy donor (HBM1) (*19*). The expression data were analyzed with Seurat (*20*) and the cells were individually annotated with SingleR (*21*) to associate the transcriptome of each cell with the gene expression profile of hematopoietic populations (*22*). The UMAP layout of these data distinguished 34 different sub-populations of hematopoietic cells **(Figure S1A)** and four differentiation pathways (erythropoiesis, granulopoiesis, dendritic maturation, and B lymphopoiesis) starting from hematopoietic stem cells (CD34+ HSC) were clearly distinguished **(Figure S1C)**. We also showed for each sub-populations the expression of classical markers (*22–25*) to validate that these data are representative of normal hematopoiesis **(Figure S1B)**.

Using the Slingshot package (*26*), we first sorted the cells along a pseudotime from the most immature (CD34+ HSC) to the most mature cells available in all the differentiation pathways with sufficient number of cells (**Supplementary table 1)**. We then computed intercellular entropy as a measure of cell-to-cell gene expression variability over a sliding window of 50 cells moving along the pseudotime with a step of 10 cells **(Figure S2)**

For erythropoiesis, 444 cells were ordered as expected: HSCs (CD34+ HSC), megakaryocyte-erythroid progenitors (CD34+ MEP), early erythroid progenitors (CD34+ ERP-Early), erythroid progenitors (CD34+ ERP), immature erythroblasts (Early-Erythroblast), and mature erythroblasts. During this differentiation pathway, we observed that intercellular entropy increases to reach a maximum at the junction between MEP and early erythroid progenitors and then falls below baseline in the mature erythroblast population **(Figure 1A)**.

**Figure 1:**
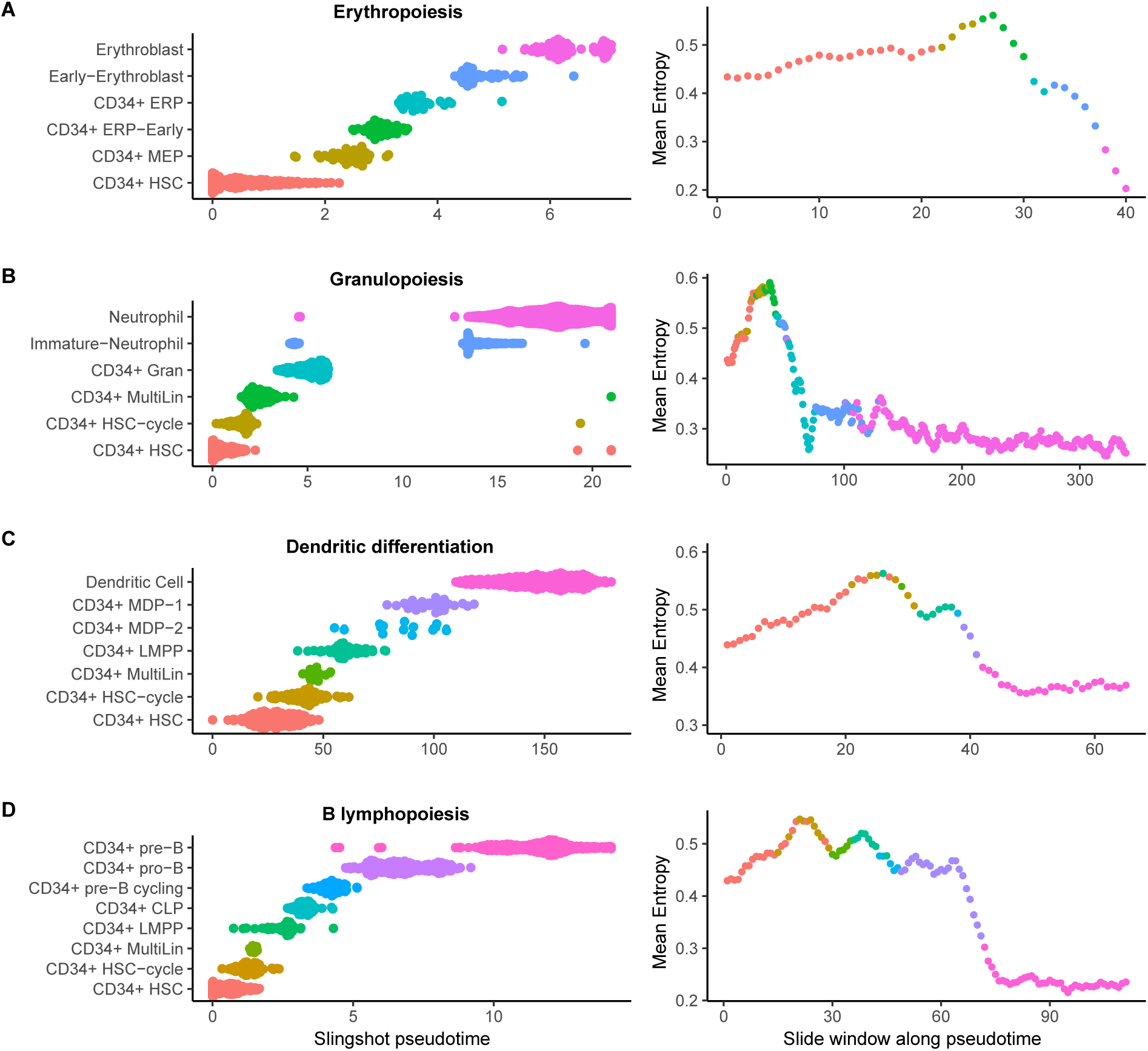
Evolution of cell-to-cell gene expression variability during the main pathways of normal hematopoietic differentiation (HBM1). Cell populations belonging to each differentiation pathway were first selected and then ordered according to the pseudotime calculated by Slingshot. The average intercellular entropy of all genes was then calculated on a sliding window of 50 cells which moves across the pseudotime with a step of 10 cells (the color of each point on the graph correspond to the nature of the first cell in the corresponding sliding window). **A)** Erythropoiesis **B)** Granulopoiesis **C)** Dendritic differentiation **D)** B Lymphopoiesis

For granulopoiesis 3440 cells were ordered as follow: HSCs, cycling HSCs (CD34+ HSC-cycle), multilineage progenitors (CD34+ MultiLin), granulocytic progenitors (CD34+ Gran), immature neutrophils and finally mature neutrophils. The measurement of intercellular entropy revealed a peak occurring at the multilineage progenitors step followed by a decrease to a minimum at the latest stages of differentiation. **(Figure 1B)**.

Regarding dendritic differentiation, 699 cells were ordered as follow: HSCs, cycling HSCs, multilineage progenitors, lympho-myeloid progenitors (CD34+ LMPP), mono-dendritic progenitors type 2 (CD34+ MDP-2), mono-dendritic progenitors type 1 (CD34+ MDP-1), and finally mature dendritic cells (Dendritic cell). During dendritic cell differentiation, intercellular entropy increased to a maximum at the junction between the cycling HSC populations and multilineage progenitors. Then, intercellular entropy decreased to a minimum in mature dendritic cells. **(Figure 1C)**.

For B lymphopoiesis we focused the analysis on the sub-populations which differentiate in the bone marrow, excluding cells which mature in lymph nodes and home back to the bone marrow (follicular and plasma cells). The 1161 cells were ordered along the pseudotime as follow and as previously descrided^22^: HSCs, cycling HSCs, multilineage progenitors, lympho-myeloid progenitors, common lymphoid progenitors (CD34+ CLP), cycling pre-B progenitors (CD34+ pre-B cycling), pro-B progenitors (CD34+ pro-B), and pre-B progenitors (CD34+ pre-B). During B lymphocyte maturation, intercellular entropy increased to reach its peak at the level of cycling HSC, then a second peak was observed at the junction between multilineage progenitors, and lympho-myeloid progenitors (CD34+ LMPP). Intercellular entropy then decreased to a minimum in the pre-B cells. **(Figure 1D)**.

Importantly, we confirmed the existence of peaks of intercellular entropy on another available scRNA-Seq dataset established on 24088 cells obtained from another normal bone marrow (HBM2) (*24*) **(Figure S3, Supplementary table 1)**

Altogether, these analyses indicate that during normal human hematopoiesis, a transient increase of cell-to-cell gene expression variability is a common feature of all of the major hematopoietic differentiation pathways.

### Variation in entropy and variation in expression highlight different set of genes

Having observed that stochastic gene expression follows the same dynamic in all the differentiation pathways, we wanted to understand what is driving this intercellular entropy peak. For each gene, we defined “delta-entropy” as the difference between maximum and minimum intercellular entropy during a given differentiation pathway, and “delta-expression” as the difference between maximum and minimum mean gene expression during differentiation **(Figure S4)**.

Intuitively, we could hypothesize that the genes with a high level of delta-expression might be those with the highest delta-entropy. Indeed, we observed a significant correlation (p < 2.10^−16^) with a moderate intensity (r < 0.34) between delta-expression and delta-entropy for each differentiation pathway **(Figure 2A).** Of note, this correlation was markedly reduced for some genes, which prompted us to analyze further the set of genes with the highest delta-entropy or delta variation. In each differentiation pathway, we selected the 20 genes with the highest delta-entropy (20-entropy) and the 20 genes with the highest delta-expression (20-expression). While most of the genes in the 20-entropy lists are specific to each differentiation pathways, the 20-expression lists were highly similar, with 14 genes common to all lineages (Common 20-expression genes) **(Figure2B, Figure S5)**. Importantly, the same observations were made on the second dataset studied previously and highlight the same specific lists of genes **(Figure S6, S7).**

**Figure 2:**
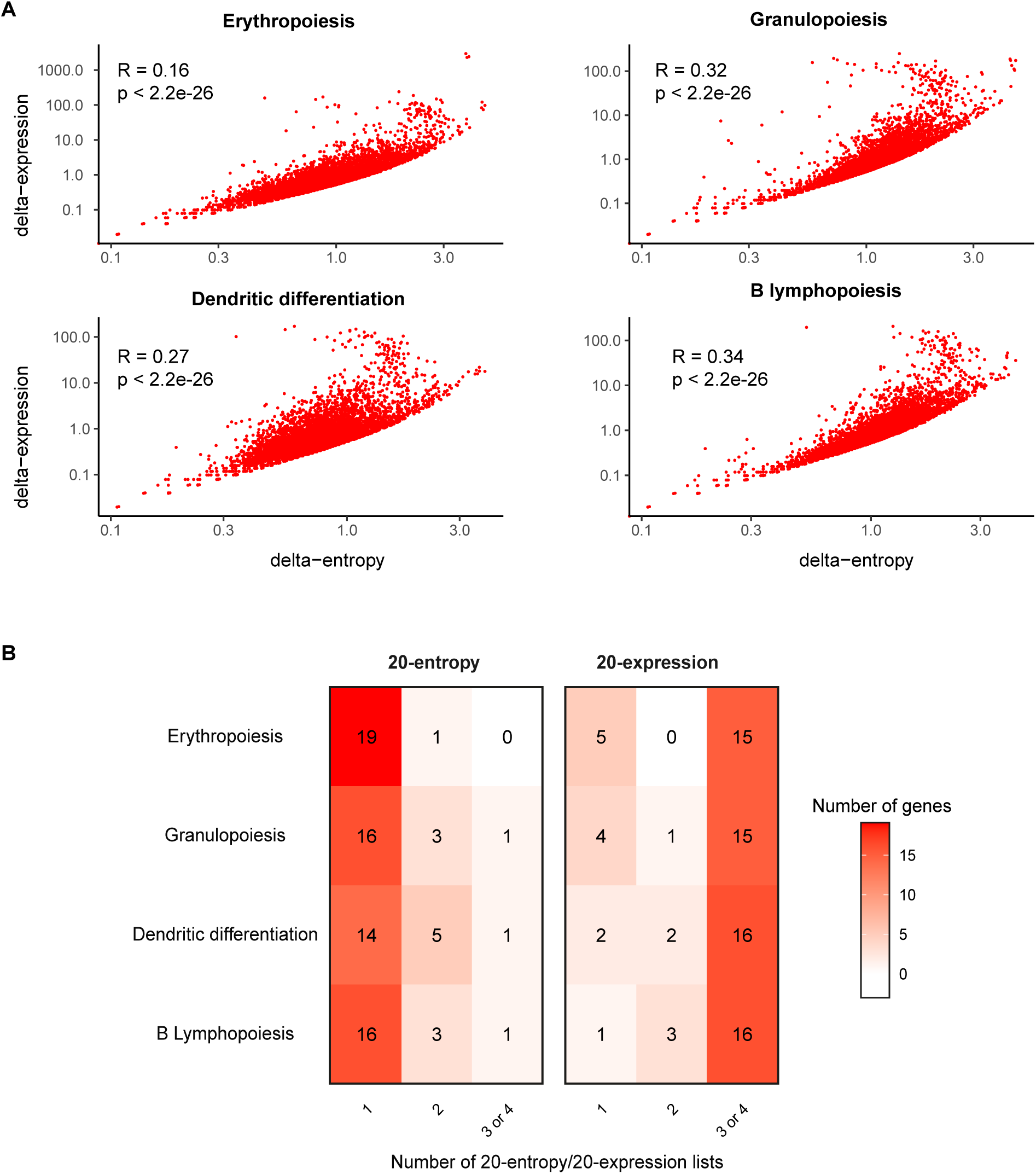
delta-entropic and delta-expressed genes along hematopoietic differentiation (HBM1) **A)** For each gene (red dots on the graphs), delta-expression is represented as a function of delta-entropy (logarithmic scale), in the 4 different hematopoiesis differentiation pathways. **B)** Overlay between the different lists. Among the 20 genes that are the most delta-entropic within the erythropoietic pathway, only 1 was also appearing in the most delta-entropic in another differentiation pathway. On the contrary, among the 20 genes with the highest delta-expression in the granulopoiesis pathway, 15 were also appearing in the 20-expression lists in at least two other differentiation pathways.

Altogether, these analyses show that during normal hematopoiesis, the highest delta-entropic genes are specific of each hematopoietic differentiation pathways, whereas most of the highest delta-expression genes are common to all differentiation pathways.

### Different functions for delta-entropic and delta-expressed genes

We used STRING database (*27*) to analyze the interaction network and functional GO enrichment of the 20-entropy gene lists and Common 20-expression genes. For each gene lists, we showed strong functional interactions between genes **(Figure3A).** Moreover, the functional GO enrichment (Biological Process and Molecular Function) in each gene list highlighted specific functions and processes of the corresponding differentiation pathway (e.g oxygen transport for erythropoiesis) **(Figure 3B)**.

**Figure 3:**
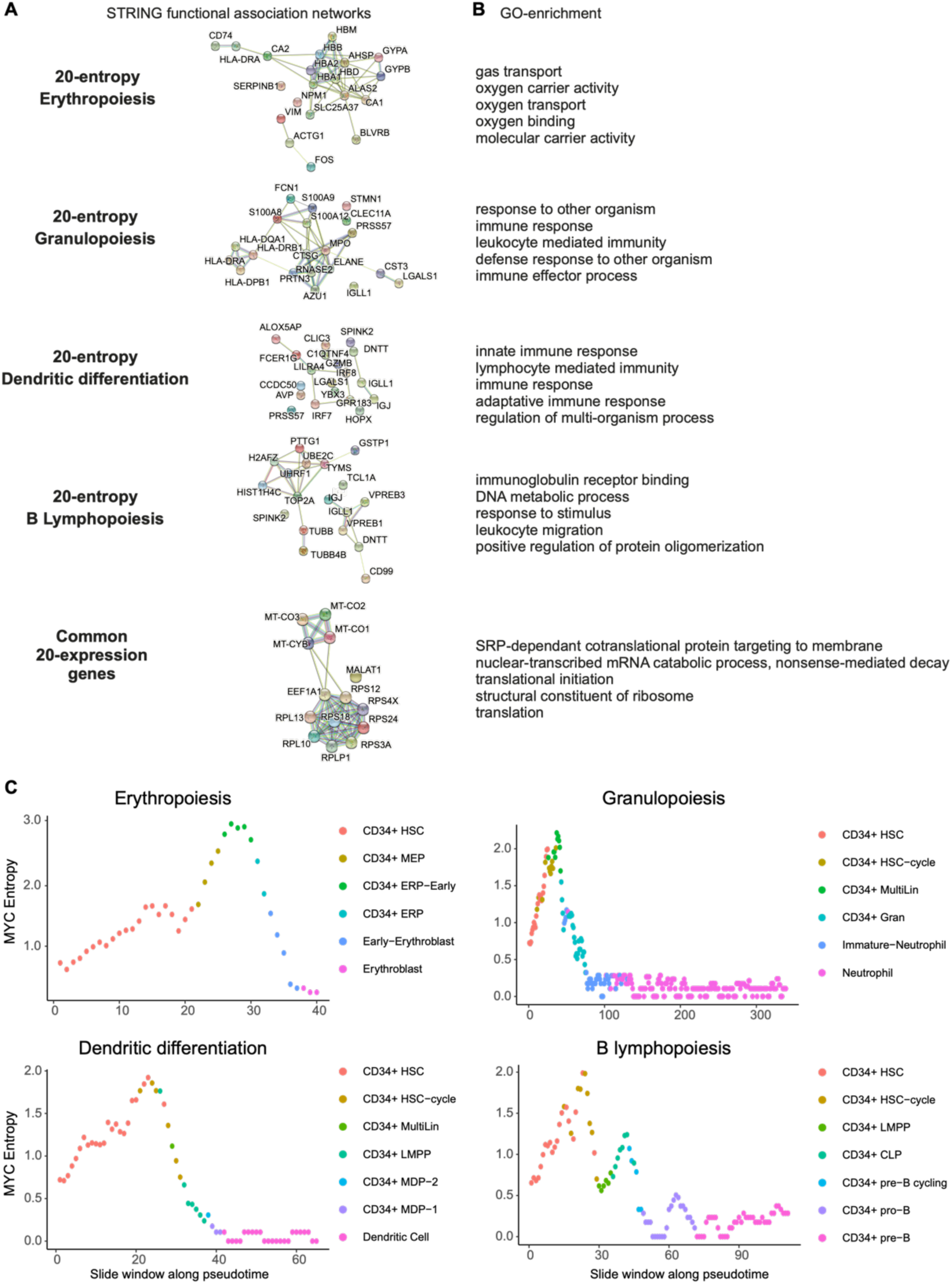
Functional association network and functional enrichment studies of 20-entropy and 20-expression gene lists. Analysis of the interaction networks **(A)** and GO functional enrichment **(B)** of the 20-entropy gene lists and Common 20-expression genes with STRING algorithm. For each pathway only the first five GO-term with a false discovery rate (FDR) lower than 0.05 were represented. **(C)** Cell-to-cell MYC expression variability during the main pathways of normal hematopoietic differentiation (HBM1).

More precisely, the 20-entropy list for erythropoiesis included 9 genes involved in hemoglobin synthesis (*HBA1, HBA2, HBB, HBD, HBM, BLVRB, ALAS2, SLC25A37, AHSP*), and 4 genes involved in other important erythropoietic process (*CA1, CA2, GYPA, GYPB*). Similarly, the 20-entropy list for granulopoiesis included alarmin genes (*S100A8, S100A9, S100A12*), genes encoding antibacterial and antiviral proteins (*AZU1, MPO, PRTN3, ELANE, CTSG, CST3, RNASE2*), antigen presenting molecules (*HLA-DPB1, HLA-DQA1, HLA-DRB1*), and lectins *(FCN1, CLEC11A*). For dendritic maturation, the list contains genes regulating the response to interferon and which are markers of mature dendritic populations and progenitors (*IRF7, IRF8*), genes playing a role in the inflammatory response (*ALOX5AP*), and innate immunity (*GZMB, C1QTNF4*). For B lymphopoiesis, the 20-entropy list comprised critical genes of B cell development such as pre-BCR formation (*VPREB1, VPREB3*), and immunoglobulin light chains (*IGLC2, IGLC3*). Interestingly, the Common 20-expression genes encode for the protein translation machinery (*RPS4X, RPL13, RPLP1, RPL10, RPS24, RPS12, RPS18, RPS3A, EEF1A*), and the mitochondrial respiratory chain (*MT-CO1, MT-CO2, MT-CO3, MT-CYB*). The long non-coding RNA (MALAT1) (*28*) was also on the Common 20-expression list.

Again, very similar results were obtained on the second dataset of normal bone marrow (HBM2).

Altogether, these analyses show that the highest delta-entropic genes are not only specific of each hematopoietic differentiation pathways but are also known to play a major role in specific functions and processes of these pathways. On the contrary most of the highest delta-expressed genes are common to all differentiation pathways, encoding proteins essential for cell survival and proliferation.

Among all the genes, those encoding transcription factors are especially important given their ability to influence the expression of a large number of genes. We conducted an analysis of the variation of the intercellular entropy of the genes identified as bona fide transcription factors. For each differentiation pathway, we selected transcription factors that belonged to the top 1000 delta-entropic genes **(Figures S8, S9, S10, S11).** We showed that transcription factors tend to have a peak of cell-to-cell gene expression variability along all 4 differentiation pathways. Moreover, MYC intercellular entropy seemed to be consistent with the mean intercellular entropy of all genes in the four differentiation pathways. **(Figure 3C).**

### A peak in cell-to-cell gene expression variability is also observed during hematopoiesis in healthy elderly subjects and SF3B1-mutated MDS

In order to further assess the role of cell-to-cell gene expression variability in pathological hematopoiesis differentiation, we reasoned that myelodysplastic syndromes (MDS) would be an interesting model. Indeed, low-risk MDS are characterized by impaired differentiation and excessive cell death of progenitors, leading to peripheral cytopenias (*29, 30*) Accordingly, we hypothesized that the analysis of the variations of cell-to-cell gene expression during differentiation could identify differences between MDS and controls. Especially for this study, we obtained and used CD34+ HSPCs from two healthy elderly controls (Ctrl1, Ctrl3) and two SF3B1 mutated MDS patients at diagnosis (MDS2, MDS4) **(Supplementary table 2).** We generated transcriptomic data of 12689 HSPC distributed over our 4 samples **(Supplementary tables 3).** In order to avoid bias due to differences in cell number in the late stages of differentiation, we focused our experiments on the CD34+ HSPC compartment, at the root of all differentiation sequences **(Figure S12A).**

We generated a reference map of all cells from the 4 samples with the integration method implemented in Seurat (*24*). The resulting UMAP allowed us to distinguish 21 different cell subtypes that are organized according to the major hematopoietic differentiation pathways (erythropoiesis, granulopoiesis, dendritic differentiation and B lymphopoiesis) **(Figure S12B).** The specific markers of the sub-populations distinguished by SingleR were consistent with previous studies of the bone marrow HSPC compartment (*23, 25, 31*) **(Figure S12C)**.

In order to compare intercellular entropy variations between MDS patients and controls, we used the integrated gene-cell matrix of the 4 samples to calculate a common pseudotime. For each differentiation pathway, we performed a subsampling in order to analyze the same number of cells in each cell type for each patient. Intercellular entropy was then computed for each patient over a sliding window of 50 cells advancing with a step of 10 on the common pseudotime.

Applying this method, a peak of intercellular entropy was observed in healthy subjects and MDS for all routes of differentiation **(Figure 4A)**

**Figure 4:**
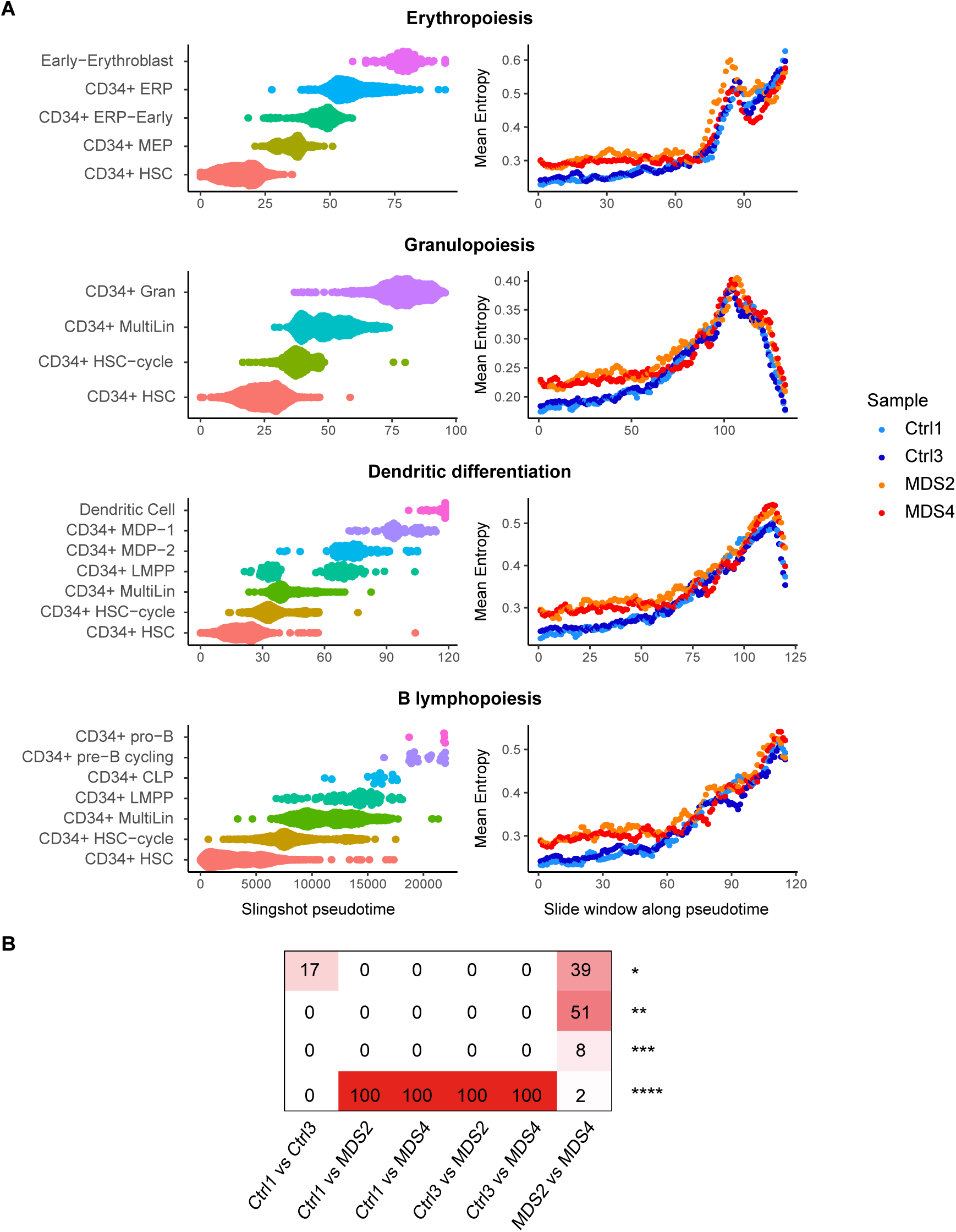
Evolution of cell-to-cell gene expression variability during hematopoiesis in elderly subjects and SF3B1-mutated MDS. **A)** For each differentiation pathway, a common pseudotime was calculated on the integrated gene cell matrix of the 4 samples. A sub-sampling was performed to have the same number of cells in each cell type per sample. The average intercellular entropy of all genes was then calculated individually for each patient on a sliding window of 50 cells advancing with a step of 10 cells on the common pseudotime. **B)** Intercellular entropy of all genes was calculated on a subsample of 700 HSCs of healthy elderly patients and SF3B1-mutated MDS. A wilcoxon assay was used to compare mean intercellular entropy between samples. This was repeated 100 times. Shown is the number of time the resulting test gave a certain level of p-value: * p<0.05, ** p<0.01, *** p<0.001, **** p <0.0001. In 100 % of the subsamples, the difference between Control and MDS patients, the mean intercellular entropy was very highly significant (p <0.0001).

For granulopoiesis, dendritic differentiation, and B lymphopoiesis, intercellular entropy peaked in the population of multilineage progenitors and then declined in more mature populations **(Figure S13, S14, S15).** For erythropoiesis, no decrease was observed, probably because of the lack of sufficient numbers of mature erythroid cells due to preanalytical steps (CD34 cell sorting) **(Figure S16).**

When observing the pattern of cell-to-cell gene expression variability in healthy elderly and MDS patients, we noted that intercellular entropy of the cells at the root of all differentiation pathways was higher in MDS than in healthy elderly subjects **(Figure 4A).** To confirm this observation, we performed 100 random subsampling of 700 HSCs in each sample and then computed the mean intercellular entropy of all genes. No difference was found between the mean intercellular entropy of HSCs of the two healthy elderly subjects. However, the HSC intercellular entropy of MDS patients was significantly higher than that of healthy elderly subjects **(Figure 4B).**

These data suggest that cell-to-cell gene expression variability is increased in MDS HSCs as compared to healthy age-matched control.

## Discussion

Differentiation can be defined as the progressive acquisition of phenotypic differences, which enables the production of highly specialized mature cells from a pool of stem cells. The driving forces of differentiation have been described according to two theoretical frameworks, highlighting the role of cell extrinsic stimuli either to initiate the process (instructive model) or to select random priming of differentiation (stochastic model) (see e.g. (*5, 12, 14, 32–34*) for elements of the discussion). Interestingly, the prediction of the evolution of cell-to-cell variability during differentiation is strongly different according to these models. According to the instructive model, cell-to-cell phenotypic variability should be limited and should not vary during differentiation, since all of the cells committed to a given lineage are following the same instructions. On the contrary, the stochastic model predicts that cell-to-cell variability should increase to reach a maximum at the stage where the selection occurs. With the technological outbreak of single cell RNA-seq, we can now use cell-to-cell gene expression variability as a proxy of cell-to-cell phenotypic variability, and provide new insights into the historical controversies between the two models of differentiation.

Using hematopoiesis as a model, we computed Shannon entropy as a measure of cell-to-cell gene expression variation. It is noteworthy that the use of Shannon entropy to analyze scRNA-seq data can be equivocal. Some authors measure the variability of gene expression in a single cell (what we called intracellular entropy) which was shown to be a proxy for stemness (*35, 36*). With this approach, the differentiation process is associated with a continuous decrease in entropy, because the transcriptional program is getting more and more specialized as cells progress in the differentiation. In the study presented here, as our aim was to capture cell-to-cell variability, we measured intercellular entropy to quantify the transcriptomic differences between cells at the same stage of differentiation.

The first important conclusion of this study is that a peak of entropy based on cell-to-cell gene expression variation is observed during the differentiation of hematopoietic stem cells along each lineage studied, strongly supporting the stochastic view of differentiation. This is in full agreement with Moussy et al (*37*) which demonstrates that human HSCs grown *in vitro* do harbor a phase characterized by fluctuating phenotypic behavior. However, the data presented here do not demonstrate any causal role of cell-to-cell gene expression variability in differentiation. This essential question could be addressed using pharmacological agents which have been demonstrated to modify the level of stochastic gene expression (*15, 18*).

Alternatively, a recent paper has reported the results of a screen identifying genes modifying cell-to-cell variability in a melanoma cell line (*38*): assessing how the knockout of these genes impacts the shape of the intercellular entropy in hematopoietic differentiation and the outcome of hematopoiesis in animal models could be very interesting.

Of note, many biological mechanisms can buffer stochastic gene expression, and reduce the phenotypic impact of the differences observed at the mRNA level(*39–42*). Notably, the half-life of proteins being significantly longer, it could buffer the variability observed at the mRNA level (*43*). Solid single cell proteomic data would represent an important step to assess the extent to which what is measurable at the mRNA level might be a good proxy of phenotypic variations.

Even if we observed a statistically significant correlation between variation in gene expression level and variation in intercellular entropy, the magnitude of this correlation remained low.

The genes with the strongest intercellular entropy variation along a given differentiation pathway were enriched in genes known to be associated with this particular lineage. By contrast, the genes with the strongest variation in mean expression level were shared by all differentiation pathways and contained genes essential for cell survival by encoding proteins involved in ribosome biogenesis, protein translation, or mitochondrial respiration. These genes are commonly described as essential in whole genome knock out screens (*44, 45*) so we can hypothesize that stochastic variations in their expression is not compatible with cell survival, explaining why we do not measure high entropy variation for these genes. The expression of these genes could be less noisy because of redundancy or other noise buffering systems (*1*). By contrast, the most highly entropic genes were very specific to each differentiation pathways. Strikingly, these specific lists comprised precisely genes already described in the differentiation pathway considered, including specific transcription factors, which enforces the biological signification of this observation.

Given the preeminent impact of differentiation abnormalities in myelodysplastic syndromes (MDS), we also assessed cell-to-cell gene expression variability in the bone marrow of two low-risk MDS patients. In this pathological hematopoiesis, the shape of the intercellular entropy variation during differentiation was conserved, but we observed a significant increase in entropy in the stem cell compartment in comparison to age matched-control samples. These data suggest that a cell-to-cell gene expression variability in the HSC compartment in MDS could be an interesting feature of this pre-leukemic process. Further studies are warranted to explore to what extent the increase of cell-to-cell variability in the stem cell compartment is contributing to MDS pathophysiology. Altogether our study is in full agreement with the wealth of recent studies that have highlighted the intensity and importance of gene expression stochasticity in all systems examined to date (*46–50*). More generally, this study highlights the need to consider the role of stochastic gene expression in complex physiological processes and pathologies such as cancer (*51, 52*) paving the way for possible noise-based therapies through epigenetic regulation (*53*).

## Patients and Methods

### Human samples

Patients Ctrl1 (female 68 years old) and Ctrl3 (male, 74 years old) were healthy elderly subjects with normal blood counts and not taking any medication that may affect hematopoiesis (chemotherapy or immunosuppressants). All patients provided written informed consent. Control samples were obtained after explanation of the study by the surgeon and the signature of a non-opposition consent. MDS2 and MDS4 patients were MDS with ring sideroblasts mutated for the SF3B1 gene and they have signed the OncoCentre Consent.

The bone marrow samples from healthy elderly patients (Ctrl1 and Ctrl3) were obtained by extraction of the bone marrow cells from the bone of the femoral head. Femoral heads were obtained after informed consent, during a surgery for hip replacement. These were cut in half and collected in a conservation medium (Hanks balanced salt solution with NaHCO3, Eurobio ™), supplemented with heparin and then transported to the laboratory at room temperature. These were scraped with a spatula, grounded in a mortar, and washed with a PBS solution supplemented with DNAse at 100 ug/ml. The mononuclear cells were then isolated on a Ficoll gradient.

The bone marrow samples from patients with myelodysplastic syndromes (MDS2, MDS4) were obtained following a bone marrow aspiration performed as part of the diagnosis of the disease. Following the sample, the mononuclear cells were isolated on a Ficoll gradient. Once isolated, mononuclear bone marrow cells were counted and resuspended in an adequate solution (IMDM, SVF 40%, DMSO 15%) and frozen in liquid nitrogen at a concentration of 20 to 30 million cells per mL. The clinical and biological characteristics of the patients are summarized in Supplementary table 2.

### Genomics studies

Bone marrow mononuclear cells were purified on Ficoll gradient and pellets were processed for nucleic acid extraction using a DNA/RNA Kit (Qiagen). Genomic DNA was studied by high throughput sequencing of 45 genes recurrently mutated in myeloid malignancies using a panel designed on Human genome hg19, and sequencing was performed on Ion PGM™ (Life Technologies) on a dedicated 318 V2 chip (*54*). Libraries were prepared using Ion AmpliSeq library kit2 384 (Life Technologies) according to the manufacturer’s instructions. Average coverage per gene was ≥500X. Reads were aligned against human genome build 19 (hg19) and analyzed for single nucleotide variant (SNV) calling with NextGENe software (SoftGenetics, Chicago, IL) and with an in-house pipeline (Polydiag, Institut Imagine, Université de Paris). We reported all clinically relevant variants with a variant allele frequency (VAF) cut-off at 2%. All the samples were also screened for ASXL1 (including c.1934dupG; p.G646WfsX12) and SRSF2 mutations by Sanger sequencing. Moreover, aligned reads from .bam files were visualized using the Integrative Genomics Viewer v2.3 from the Broad Institute (Cambridge, MA, USA). Assessment of variants implication was performed based on population databases (dbSNP and GnomAD), mutation databases (COSMIC), and predictions software (Alamut, mutation taster, OncoKB, and Cancer Genome Interpreter).

### Single cell RNA-seq

Bone marrow mononuclear cells were thawed, and dead cells were eliminated by immunomagnetic negative sorting (MACS MicroBead technology from Miltenyi Biotec™). A second CD34 positive immunomagnetic sorting was performed to isolate stem and progenitor cells. Cells were then washed with PBS containing 0.04% of BSA. Cell concentration and viability were determined microscopically with a Malassez counting chamber cell after staining with trypan blue.

Libraries were prepared with the chromium system of 10x genomics, with the Chromium Single Cell 3 ’V2 kit according to the manufacturer’s protocol (www.10xgenomics.com). The four samples (Ctrl1, Ctrl3, MDS2 and MDS4) were processed on the same chromium chip. The number of cells targeted per sample was 5000 cells. The libraries were sequenced by the Integragen company on an Illumina HiSeq4000 sequencer with a target depth of 50,000 reads per cell. Sequencing data are deposited in the Gene Expression Omnibus (GEO) with the accession code GSE169426.

### Bioinformatic analysis

#### Gene-cell expression matrices

Healthy donor total bone marrow scRNA-seq datasets used in this study were published by Granja et al (HBM1) (*19*) and Stuart et al (HBM2) (*24*).

For HBM1, gene-cell expression matrix files were downloaded from: https://www.ncbi.nlm.nih.gov/geo/query/acc.cgi?acc=GSE139369 (GSM4138872_scRNA_BMMC_D1T1.rds.gz, GSM4138873_scRNA_BMMC_D1T2.rds.gz). Since the authors removed the ribosomal and mitochondrial genes from the gene-cell matrix, the original .bam files were downloaded from https://trace.ncbi.nlm.nih.gov/Traces/sra/?run=SRR10343065 (SRR10343065/scRNA_BMMC_D1T1.bam.1) https://trace.ncbi.nlm.nih.gov/Traces/sra/?run=SRR10343066 (SRR10343066/scRNA_BMMC_D1T2.bam.1), and therefore processed using the DropEst (*55*) software. The expression of the mitochondrial and ribosomal genes thus obtained was then reincorporated into the previously filtered gene-cell matrix.

For HBM2, gene-cell expression matrix files were downloaded from https://www.ncbi.nlm.nih.gov/geo/query/acc.cgi?acc=GSM3681518 (GSM3681518_MNC_RNA_counts.tsv.gz), and https://www.ncbi.nlm.nih.gov/geo/query/acc.cgi?acc=GSM3681520 (GSM3681520_MNC_HTO_counts.tsv.gz).

For Ctrl1, Ctrl3, MDS2 and MDS4 samples, the Cell Ranger software (https://support.10xgenomics.com/single-cellgene-expression/software/pipelines/latest/what-is-cell-ranger) was used to process the raw data from Illumina sequencing and generate the gene-cell expression matrices. The reads were aligned on the GRCh38 reference genome by STAR with the ENSEMBL annotation.

#### Gene expression matrix filtering

For HBM1, we keep the same filters as the authors did (*19*).

For HBM2, we chose to keep for downstream analysis the cells which expressed between 500 and 4000 genes with a percentage of mitochondrial genes less than 15%. For Ctrl1, Ctrl2, MDS2 and MDS4, we chose to keep for downstreamn analysis the cells which expressed between 500 and 5500 genes with a percentage of mitochondrial genes less than 10%.

#### Dimensionnality reduction

The gene-cell expression matrices of each sample were normalized with SCtransform (*56*). After normalization, the gene-cell matrices were subjected to dimensionality reduction techniques such as principal component analysis (PCA) and UMAP (*57*) using Seurat (*24*). We also used the Scanpy (*58*) python package to calculate the ForceAtlas2 (FA). For HBM1 and HBM2, we chose not to use ribosomal and mitochondrial genes in order to improve the results.

#### Cell annotation

The cells were annotated individually using the SingleR(*21*) software by comparing their gene expression profile with the 34 bone marrow populations published by Hay et al (*22*).

#### Pseudotime ordering

For each sample individually, the cells were ordered from HSCs to mature cells for each differentiation pathway. A pseudotime was computed with Slingshot(*26*) with the following options : the starting cell population was always specified as CD34 + HSC; the multidimensional space specified was either the UMAP, the FA, or the PCA; the final pseudotime cell population was sometimes specified as the subpopulation corresponding to the most mature cells of the differentiation pathway studied. Thus, depending on the options chosen, several pseudotimes were calculated for each differentiation path. The pseudotime chosen for intercellular entropy analyzes being the one which allowed the cells to be ordered for each differentiation pathway in the most consistent way with our knowledge of hematopoiesis.

For Ctrl1, Ctrl3, MDS2, and MDS4, gene-cell expression matrices were integrated using Seurat (*24*) after normalization with SCtransform technique in order to compute common multidimensional spaces (PCA, UMAP, and FA). This allowed slingshot to compute a common pseudotime on the four differenciation pathways.

#### Intercellular entropy computation

For each sample, the intercellular entropy was computed on the raw counts of the filtered gene expression matrices for each differentiation pathway. We compute an intercellular Shannon entropy per gene, and displayed the mean entropy variation along the differentiation sequence. Since we average this value, it will not be affected by the number of genes which may vary as a function of time.

Because Ctrl1 and Ctrl3 cells come from sample that were mechanically dissociated, we chose not to include “dissociation genes” for the comparison of intercellular entropy between elderly subjects and SF3B1 mutated MDS. These “dissociation genes” were the common genes between two dissociation signatures previously published (*59, 60*).

### Shannon Entropy estimation

To estimate the Shannon entropy, the Best Upper Bound estimator (BUB) was computed as developed in Paninski et al (*61*). The N/m ratio is critical for the choice of the proper entropy estimator, with N being the number of cells used to compute the entropy, and m being the number of bins, here the largest number of RNA molecule found in each cell. For small N/m ratios (less than 100) the BUB was shown to outperform the Maximum Likelihood Estimator, the Miller-Madow or the Jackknifed estimator, by minimizing the maximum error. Since we are using sliding windows of 50 cells, the choice of the BUB estimator prevailed. The R script for this estimation is available on this OSF project: https://osf.io/9mcwg/

## Acknowledgements

This work was supported by the Assocation Laurette Fugain (2017) and by a grant from Institut Convergence PLASCAN (ANR-17-CONV-0002).

CD was the recipient of a PhD funding from aviesan ITMO Cancer (Plan Cancer 2014-2019).

The authors would like to thank Thomas Mercher for his insightful suggestions and careful reading of the manuscript.

---

**Figure S1:**
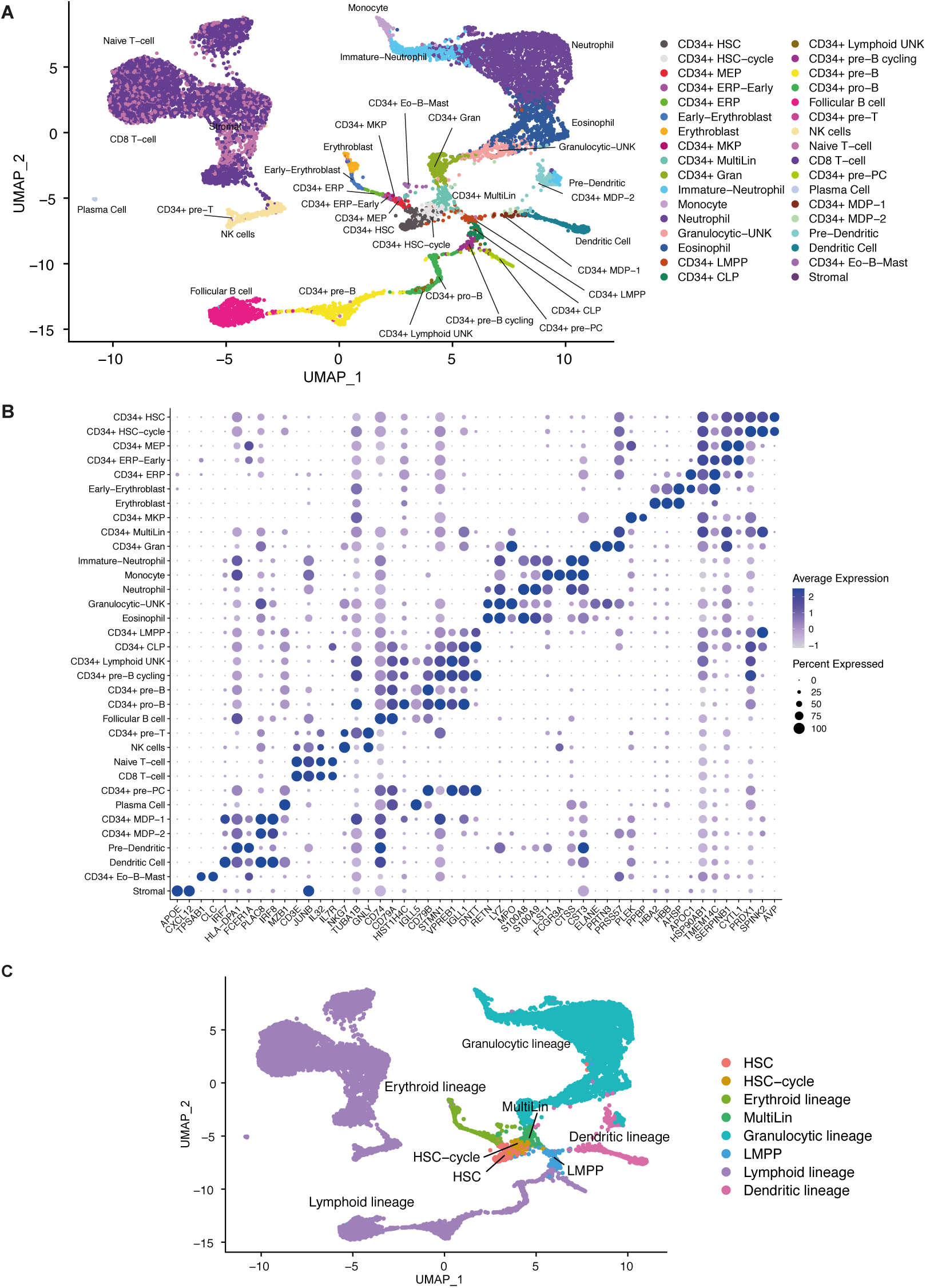
Single cell transcriptomic landscape of healthy human bone marrow (HBM1). **A)** UMAP of scRNAseq data on healthy donor bone marrow mononuclear cells published by Granja et al (*19*), 34 of the 35 populations described by Hay et al (*22*) were distinguished by SingleR. **B)** Expression values of selected marker genes for all cell sub-populations. Circle color shows mean scaled expression values and circle size represents the proportion of expressing cells per sub-populations. **C)** UMAP of Main hematopoietic differentiation pathways (erythropoiesis, granlulopoiesis, B lymphopoiesis and dendritic maturation).

**Figure S2:**
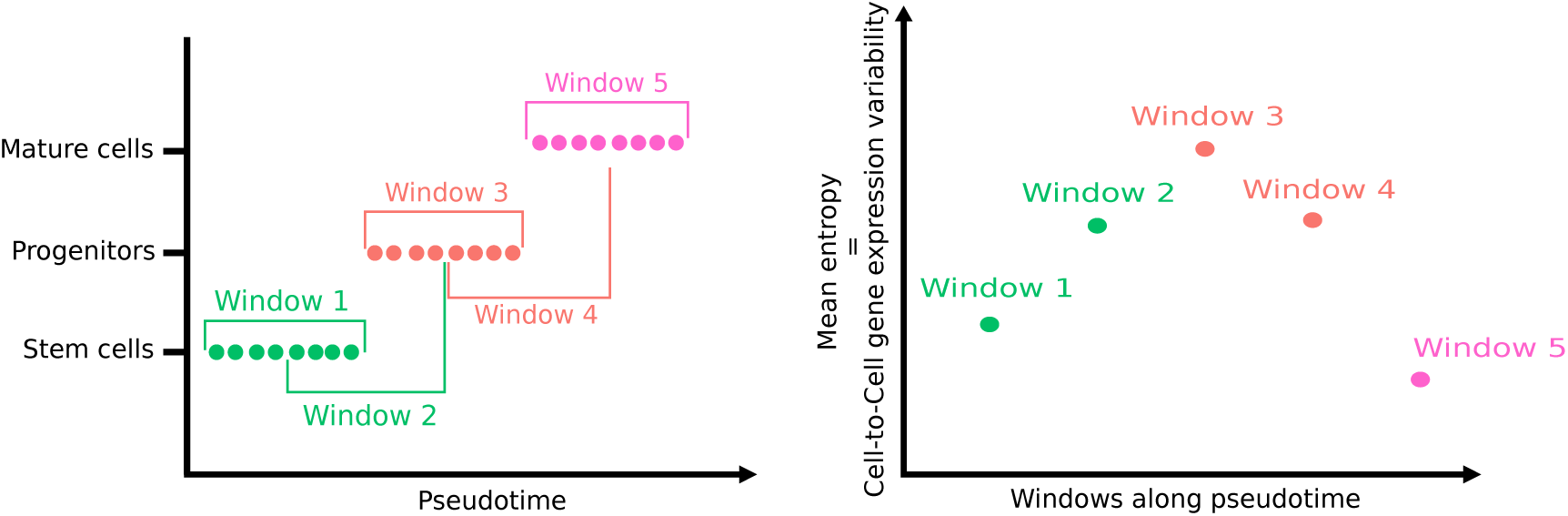
Strategy used to represent the evolution of cell-to-cell gene expression variability during differentiation. Cells are first ordered according to their position in the pseudotime which reflects their progress across the differentiation. The mean intercellular entropy of all genes was then calculated on a window of (for the example) 8 cells (window1). The window is moving across pseudotime (window2, window3…) with a step of (for the example) 4 cells, and the mean intercellular entropy is calculated for every windows. The color of each point on the graph (right panel) correspond to the nature of the first cell in the corresponding window.

**Figure S3:**
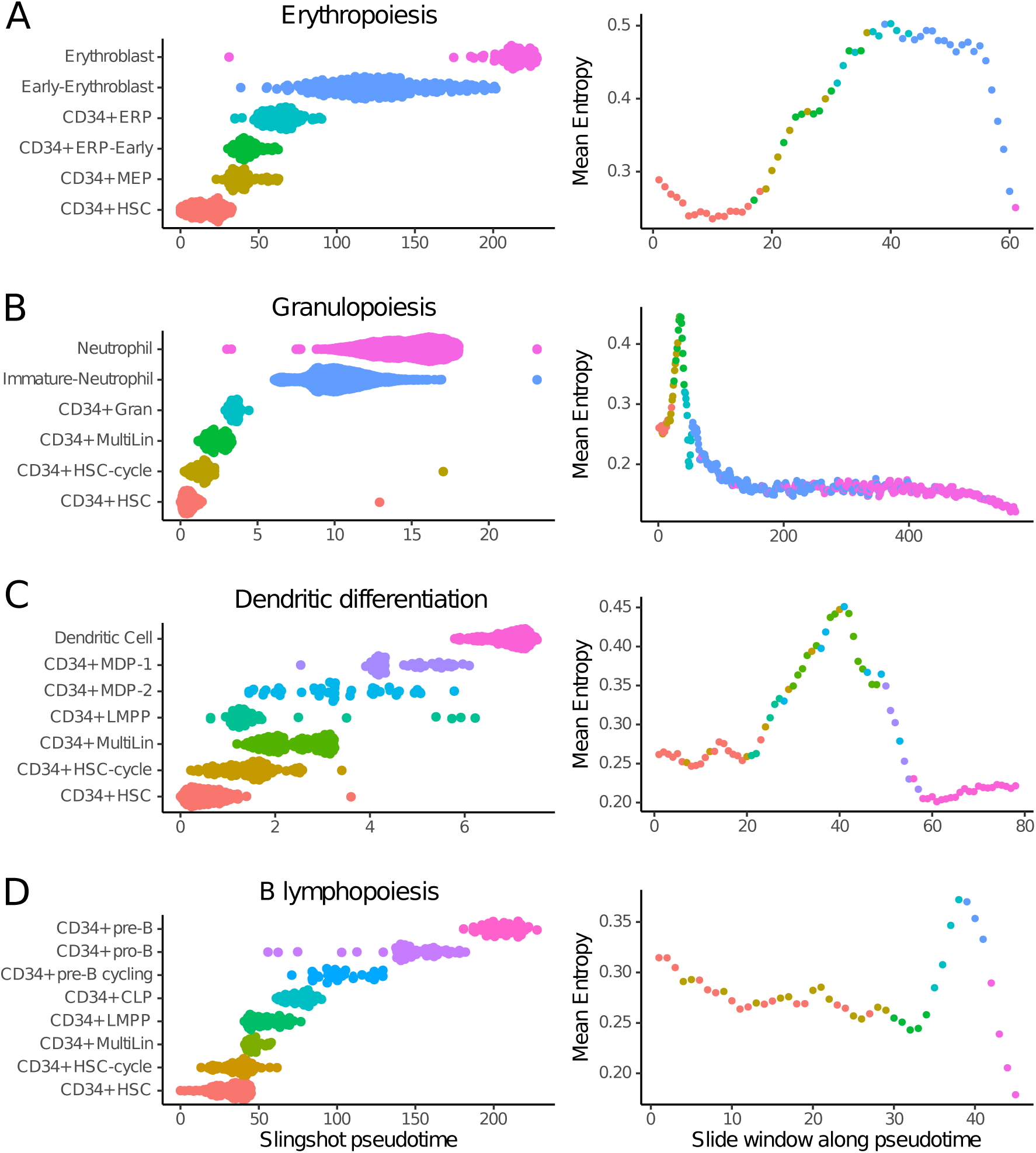
Evolution of cell-to-cell gene expression variability during the main pathways of normal hematopoietic differentiation (HBM2). Cell populations belonging to each differentiation pathway were first selected and then ordered according to the pseudotime calculated by Slingshot. The average intercellular entropy of all genes was then calculated on a sliding window of 50 cells which moves across the pseudotime with a step of 10 cells (the color of each point on the graph correspond to the nature of the first cell in the corresponding sliding window). **A)** Erythropoiesis **B)** Granulopoiesis **C)** Dendritic differentiation **D)** B Lymphopoiesis

**Supplementary table 1:** Distribution of cellular subpopulations for each differentiation pathway in HBM1 and HBM2 dataset.

HBM1
| Erythropoiesis | CD34+<br>HSC | CD34+<br>MEP | CD34+<br>ERP-Early | CD34+<br>ERP | Early-<br>Erythroblast | Erythroblast |
| --- | --- | --- | --- | --- | --- | --- |
| # cells | 204 | 39 | 45 | 23 | 54 | 79 |

| Granulopoiesis | CD34+<br>HSC | CD34+<br>HSC-cycle | CD34+<br>MultiLin | CD34+<br>Gran | Immature-<br>Neutrophil | Neutrophil |
| --- | --- | --- | --- | --- | --- | --- |
| # cells | 209 | 99 | 117 | 277 | 416 | 2322 |

| Dendritic<br>differentiation | CD34+<br>HSC | CD34+<br>HSC-cycle | CD34+<br>MultiLin | CD34+<br>LMPP | CD34+<br>MDP-2 | CD34+<br>MDP-1 | Dendritic Cell |
| --- | --- | --- | --- | --- | --- | --- | --- |
| # cells | 198 | 98 | 109 | 62 | 12 | 29 | 289 |

| B<br>Lymphopoiesis | CD34+<br>HSC | CD34+<br>HSC-cycle | CD34+<br>MultiLin | CD34+<br>LMPP | CD34+<br>CLP | CD34+<br>pre-B<br>cycling | CD34+<br>pro-B | CD34+<br>Pre-B |
| --- | --- | --- | --- | --- | --- | --- | --- | --- |
| # cells | 205 | 98 | 116 | 63 | 72 | 29 | 289 | 455 |

HBM2
| Erythropoiesis | CD34+<br>HSC | CD34+<br>MEP | CD34+<br>ERP-Early | CD34+<br>ERP | Early-<br>Erythroblast | Erythroblast |
| --- | --- | --- | --- | --- | --- | --- |
| # cells | 163 | 69 | 72 | 100 | 180 | 74 |

| Granulopoiesis | CD34+<br>HSC | CD34+<br>HSC-cycle | CD34+<br>MultiLin | CD34+<br>Gran | Immature-<br>Neutrophil | Neutrophil |
| --- | --- | --- | --- | --- | --- | --- |
| # cells | 164 | 114 | 140 | 102 | 2273 | 2956 |

| Dendritic<br>Differentiation | CD34+<br>HSC | CD34+<br>HSC-cycle | CD34+<br>MultiLin | CD34+<br>LMPP | CD34+<br>MDP-2 | CD34+<br>MDP-1 | Dendritic Cell |
| --- | --- | --- | --- | --- | --- | --- | --- |
| # cells | 164 | 114 | 140 | 42 | 37 | 56 | 272 |

| B Lymphopoiesis | CD34+<br>HSC | CD34+<br>HSC-cycle | CD34+<br>MultiLin | CD34+<br>LMPP | CD34+<br>CLP | CD34+<br>pre-B<br>cycling | CD34+<br>pro-B | CD34+<br>Pre-B |
| --- | --- | --- | --- | --- | --- | --- | --- | --- |
| # cells | 164 | 106 | 23 | 42 | 40 | 23 | 62 | 40 |

**Figure S4:**
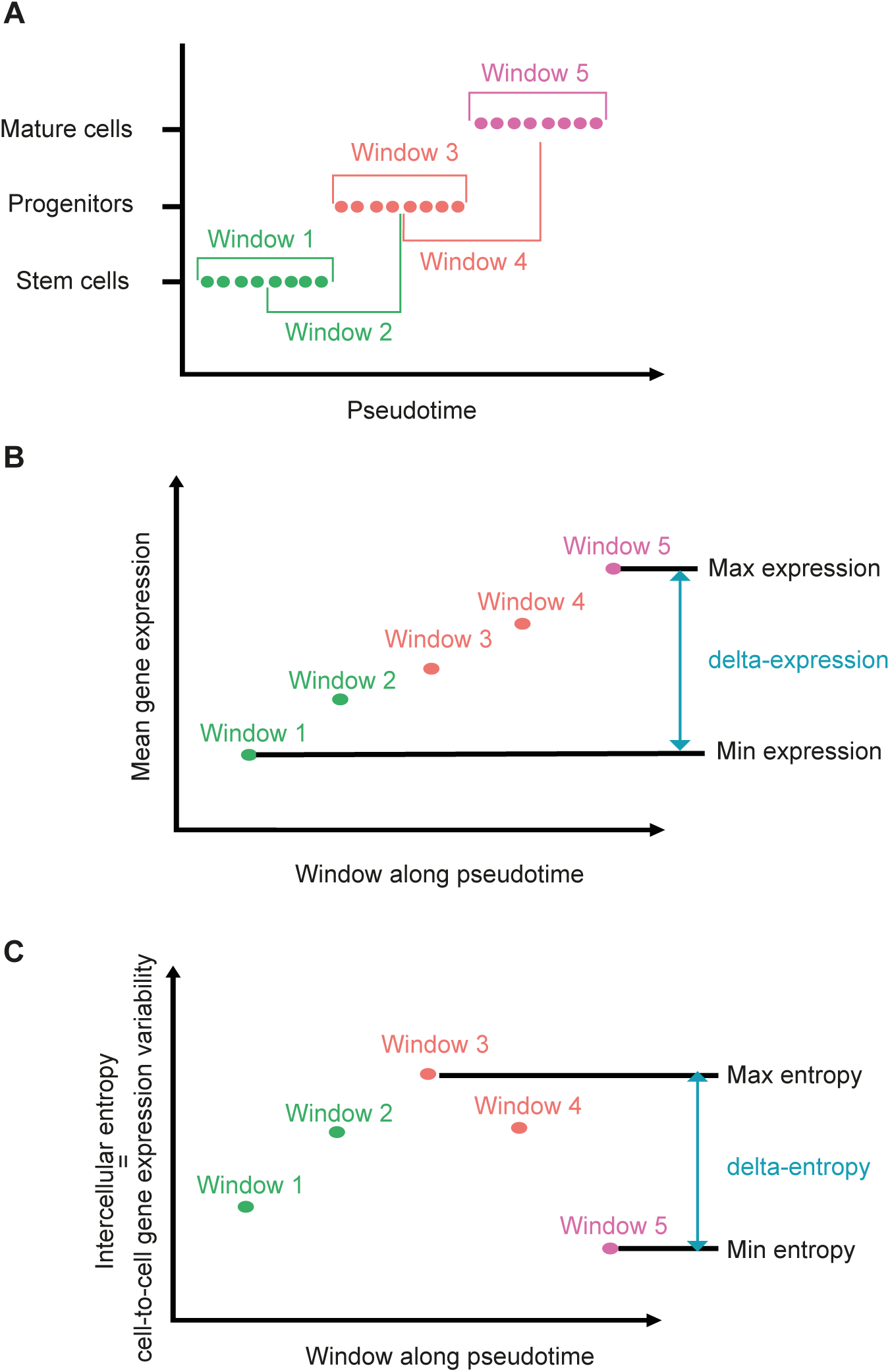
Strategy used to calculate delta-entropy and delta-expression. **(A)** Cells are first ordered according to their position in the pseudotime which reflects their progress across the differentiation. **(B)** The mean expression of a gene is then calculated for each window. For each gene, the difference between minimum and maximum expression is the delta-expression **(C)** The intercellular entropy of a gene is also calculated for each window. For each gene, the difference between minimum and maximum entropy is the delta-entropy.

**Figure S5:**
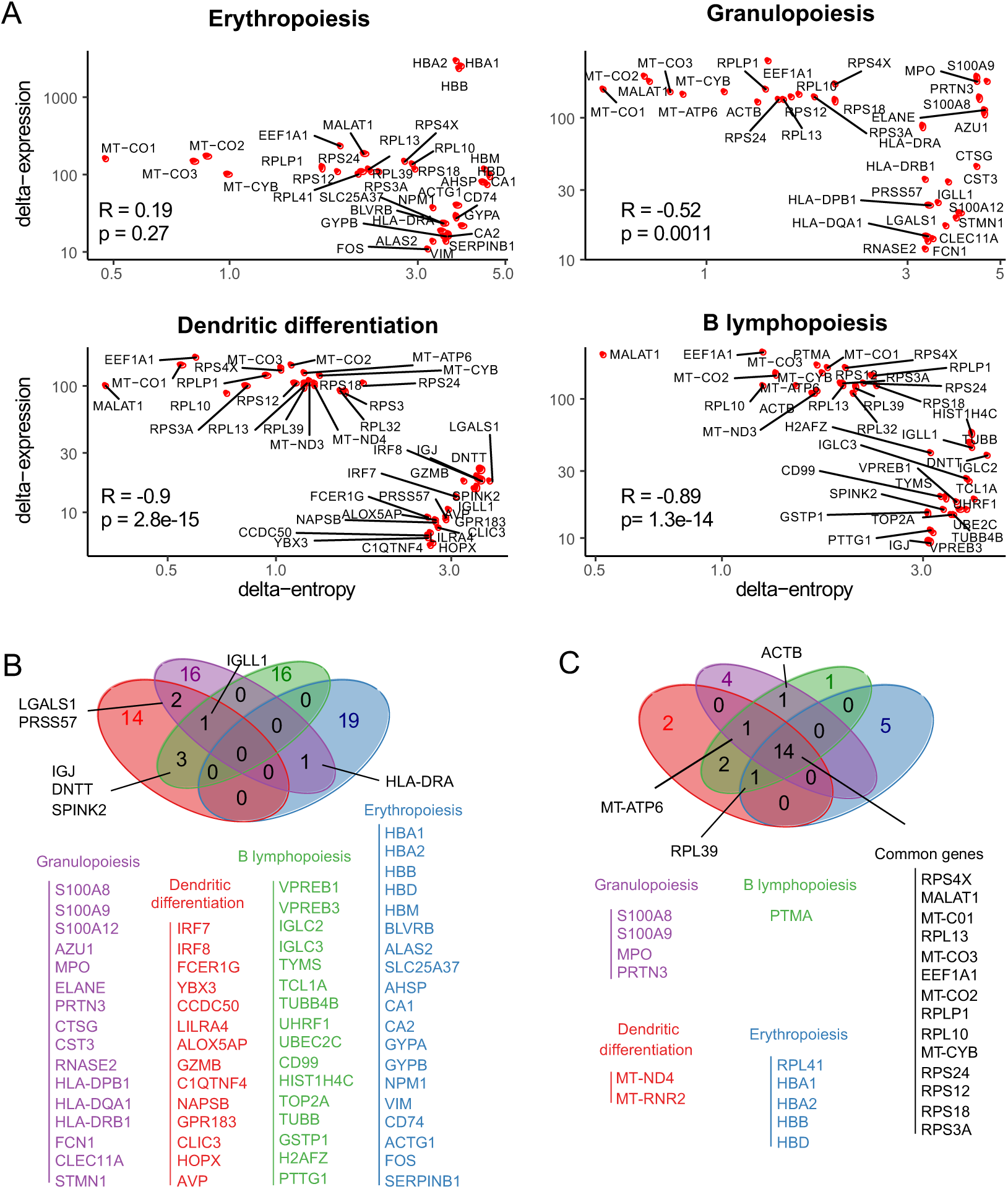
Most delta-entropic and most delta-expressed genes along hematopoietic differentiation (HBM1). **(A)** For each gene (red dots on the graphs), delta-expression is represented as a function of delta-entropy (logarithmic scale), in the 4 different hematopoiesis differentiation pathways. **B-C)** Venn diagram of the 20 most delta entropic **(B)** and 20 most delta expressed **(C)** genes during the different hematopoietic differentiation pathways.

**Figure S6:**
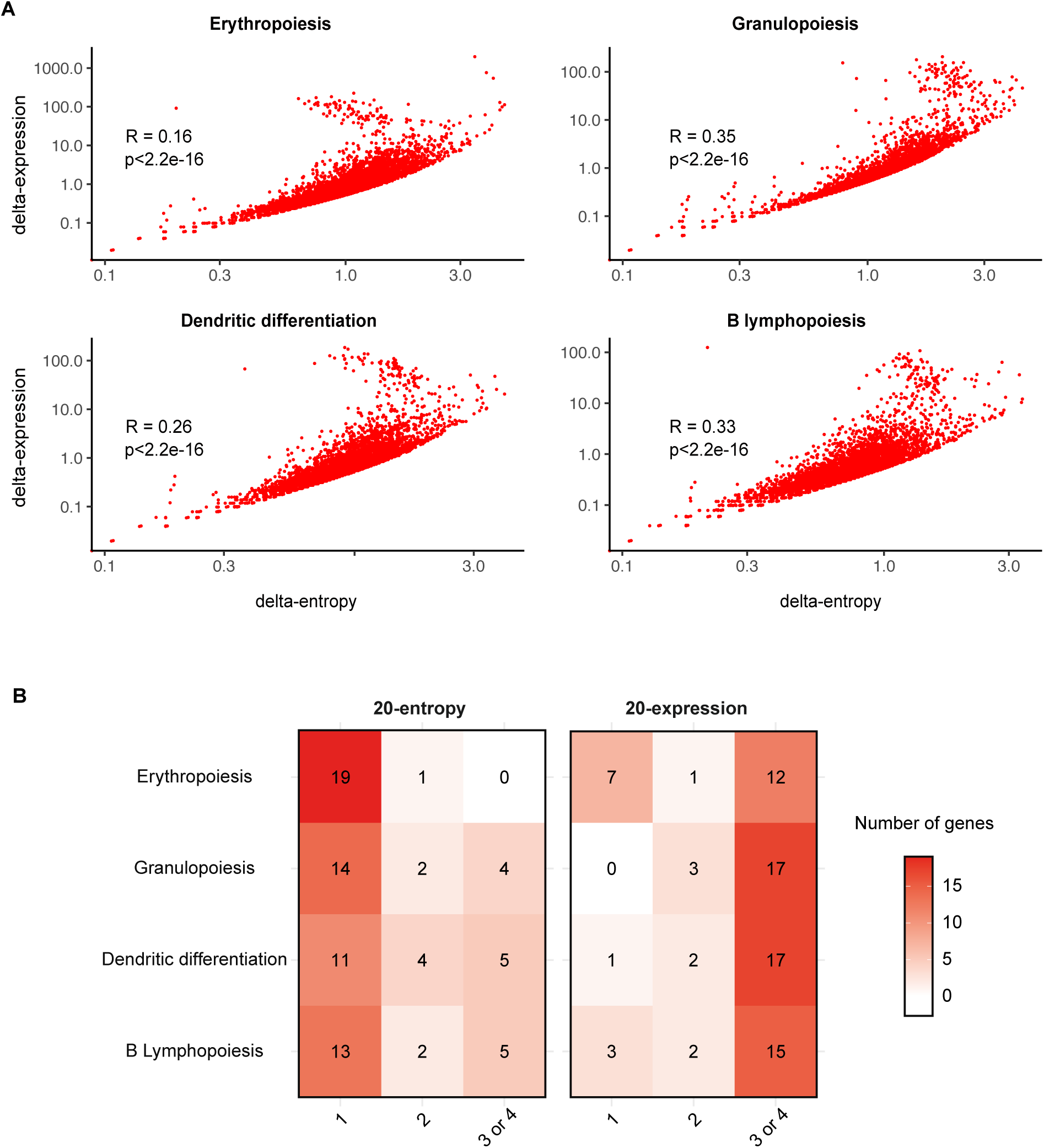
delta-entropic and delta-expressed genes along hematopoietic differentiation (HBM2) **A)** For each gene (red dots on the graphs), delta-expression is represented as a function of delta-entropy (logarithmic scale), in the 4 different hematopoiesis differentiation pathways. **B)** Overlay between the different lists. Among the 20 genes that are the most delta-entropic within the erythropoietic pathway, only 1 was also appearing in the most delta-entropic in another differentiation pathway. On the contrary, among the 20 genes with the highest delta-expression in the granulopoiesis pathway, 17 were also appearing in the 20-expression lists in at least two other differentiation pathways.

**Figure S7:**
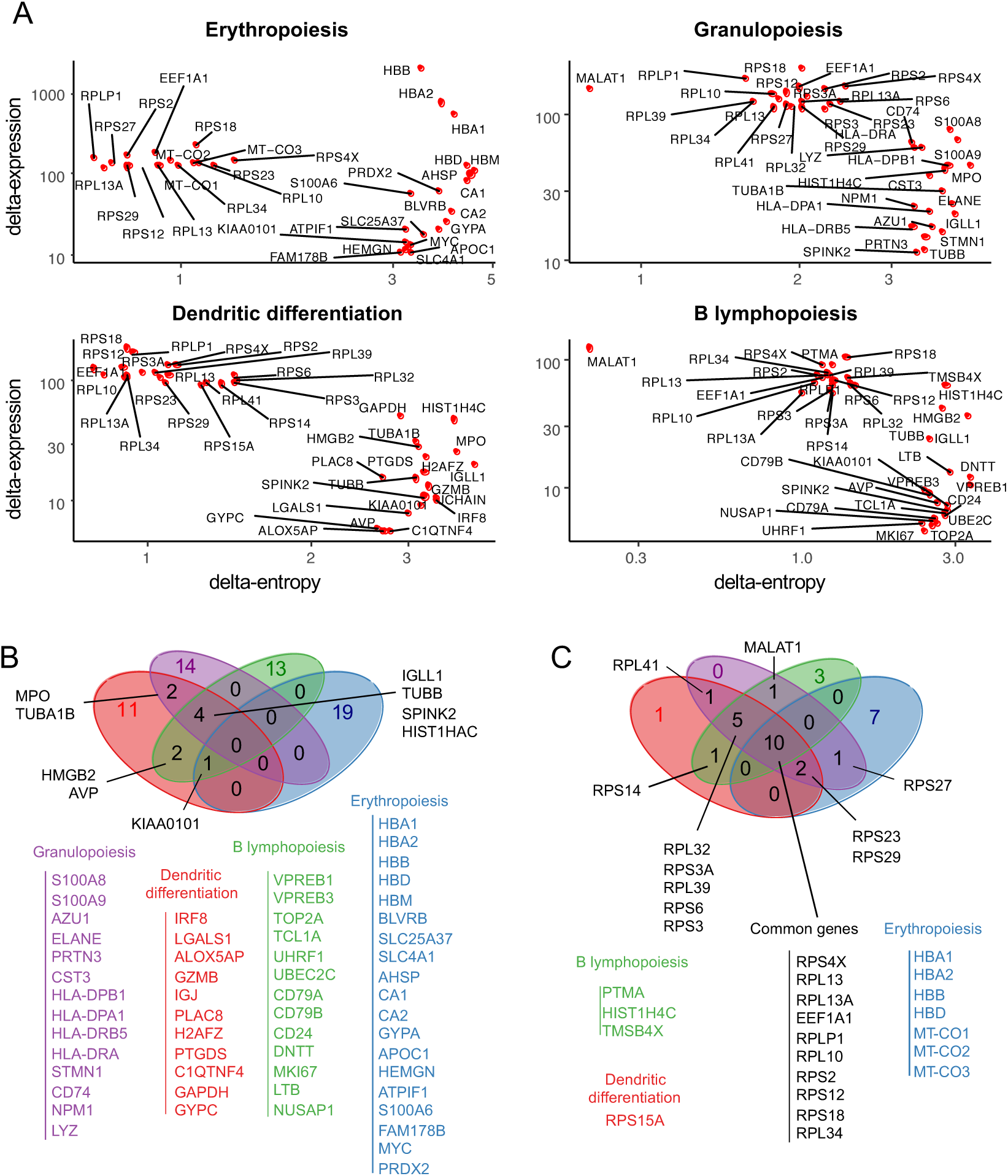
Most delta-entropic and most delta-expressed genes along hematopoietic differentiation (HBM2). **(A)** For each gene (red dots on the graphs), delta-expression is represented as a function of delta-entropy (logarithmic scale), in the 4 different hematopoiesis differentiation pathways. **B-C)** Venn diagram of the 20 most delta entropic **(B)** and 20 most delta expressed **(C)** genes during the different hematopoietic differentiation pathways.

**Figure S8:**
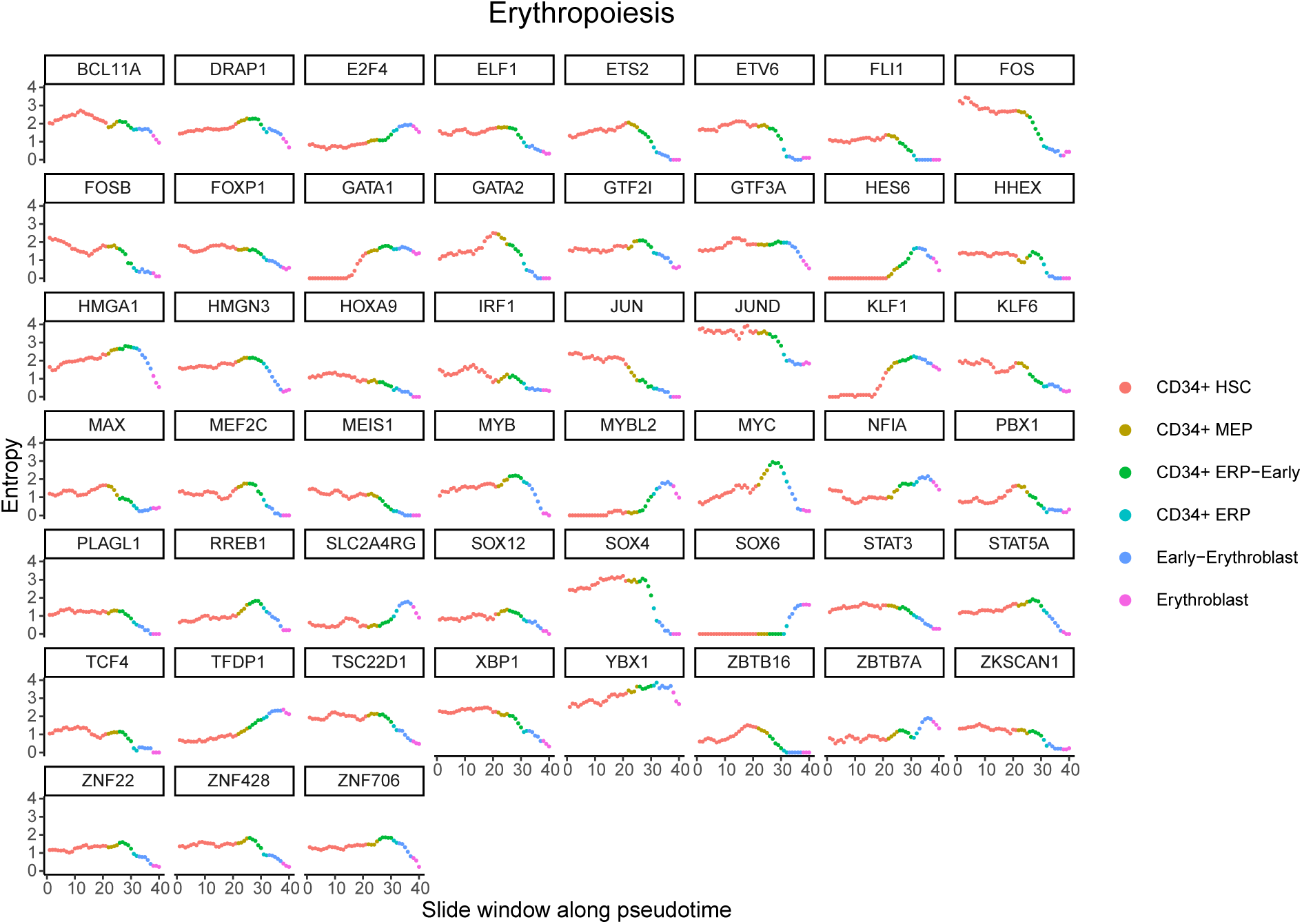
Cell-to-cell gene expression variability of transcription factors belonging to the 1000 most delta entropic genes during Erythropoiesis (HBM1). Cell populations belonging to erythropoiesis were first selected and then ordered according to the pseudotime calculated by Slingshot. The intercellular entropy of each transcription factor was then calculated on a sliding window of 50 cells which moves across the pseudotime with a step of 10 cells (the color of each point on the graph correspond to the nature of the first cell in the corresponding sliding window).

**Figure S9:**
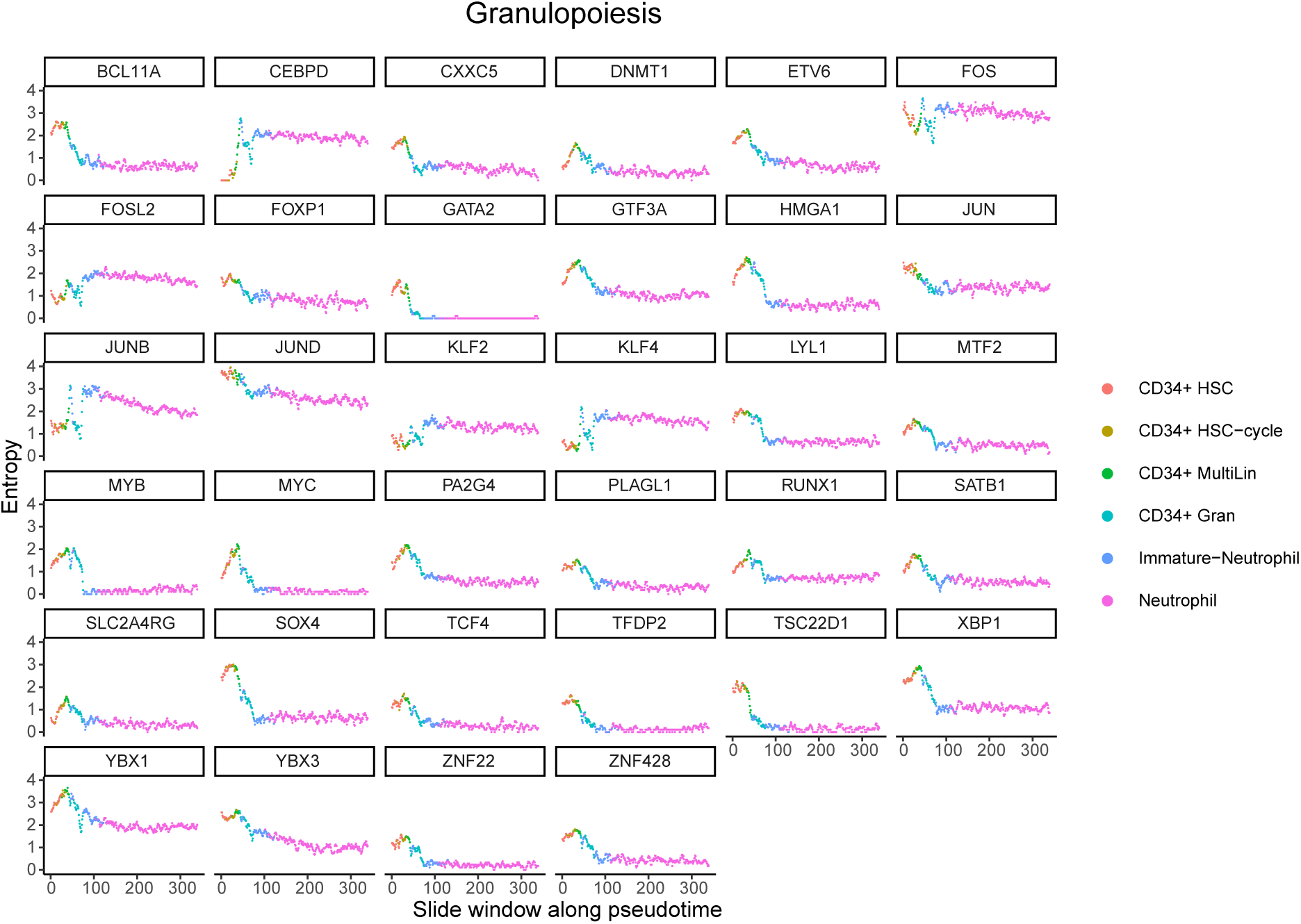
Cell-to-cell gene expression variability of transcription factors belonging to the 1000 most delta entropic genes during Granulopoiesis (HBM1). Cell populations belonging to granulopoiesis were first selected and then ordered according to the pseudotime calculated by Slingshot. The intercellular entropy of each transcription factor was then calculated on a sliding window of 50 cells which moves across the pseudotime with a step of 10 cells (the color of each point on the graph correspond to the nature of the first cell in the corresponding sliding window).

**Figure S10:**
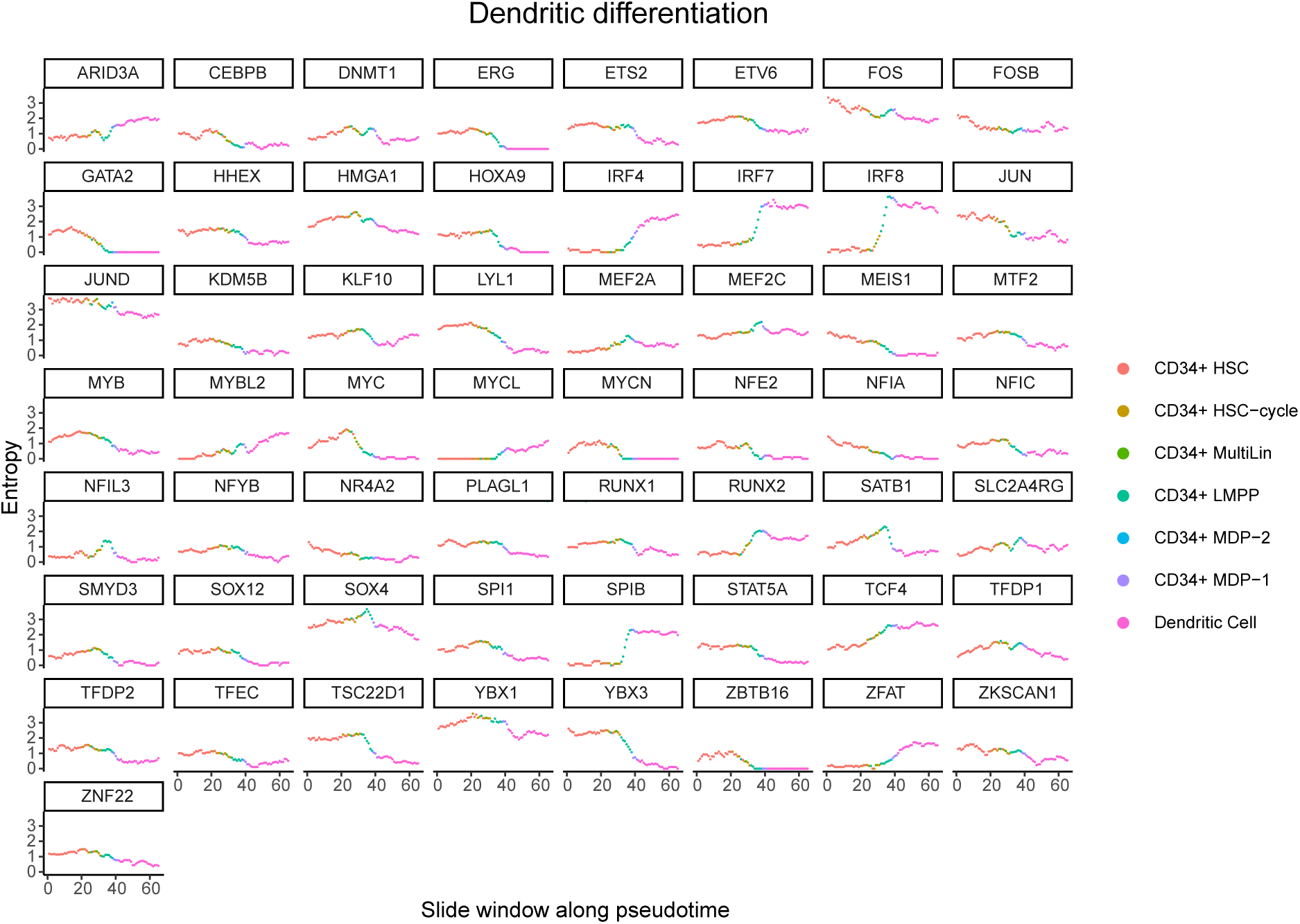
Cell-to-cell gene expression variability of transcription factors belonging to the 1000 most delta entropic genes during dendritic differentiation (HBM1). Cell populations belonging to dendritic differentiation were first selected and then ordered according to the pseudotime calculated by Slingshot. The intercellular entropy of each transcription factor was then calculated on a sliding window of 50 cells which moves across the pseudotime with a step of 10 cells (the color of each point on the graph correspond to the nature of the first cell in the corresponding sliding window).

**Figure S11:**
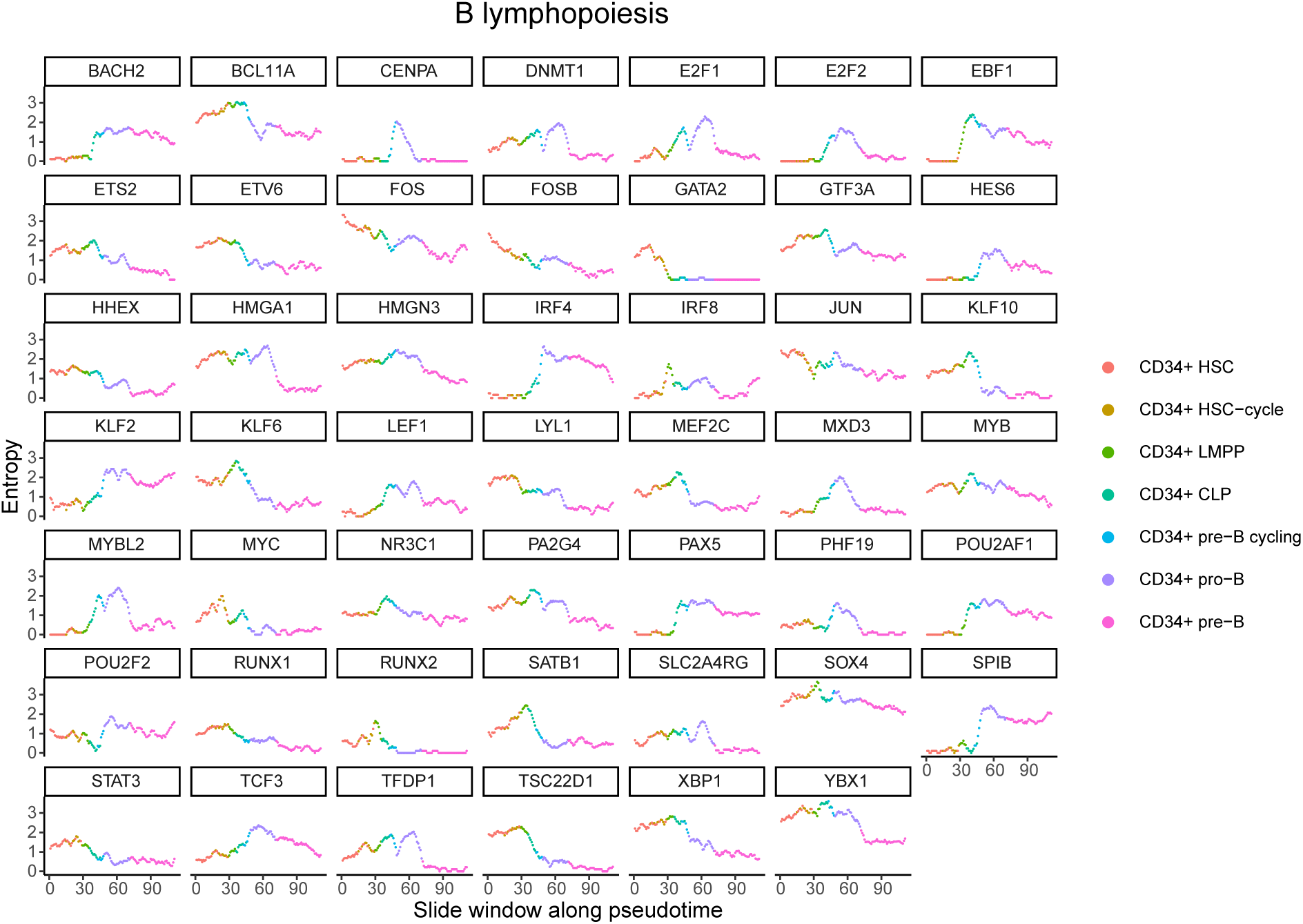
Cell-to-cell gene expression variability of transcription factors belonging to the 1000 most delta entropic genes during B lymphopoiesis (HBM1). Cell populations belonging to B lymphopoieses were first selected and then ordered according to the pseudotime calculated by Slingshot. The intercellular entropy of each transcription factor was then calculated on a sliding window of 50 cells which moves across the pseudotime with a step of 10 cells (the color of each point on the graph correspond to the nature of the first cell in the corresponding sliding window).

**Figure S12:**
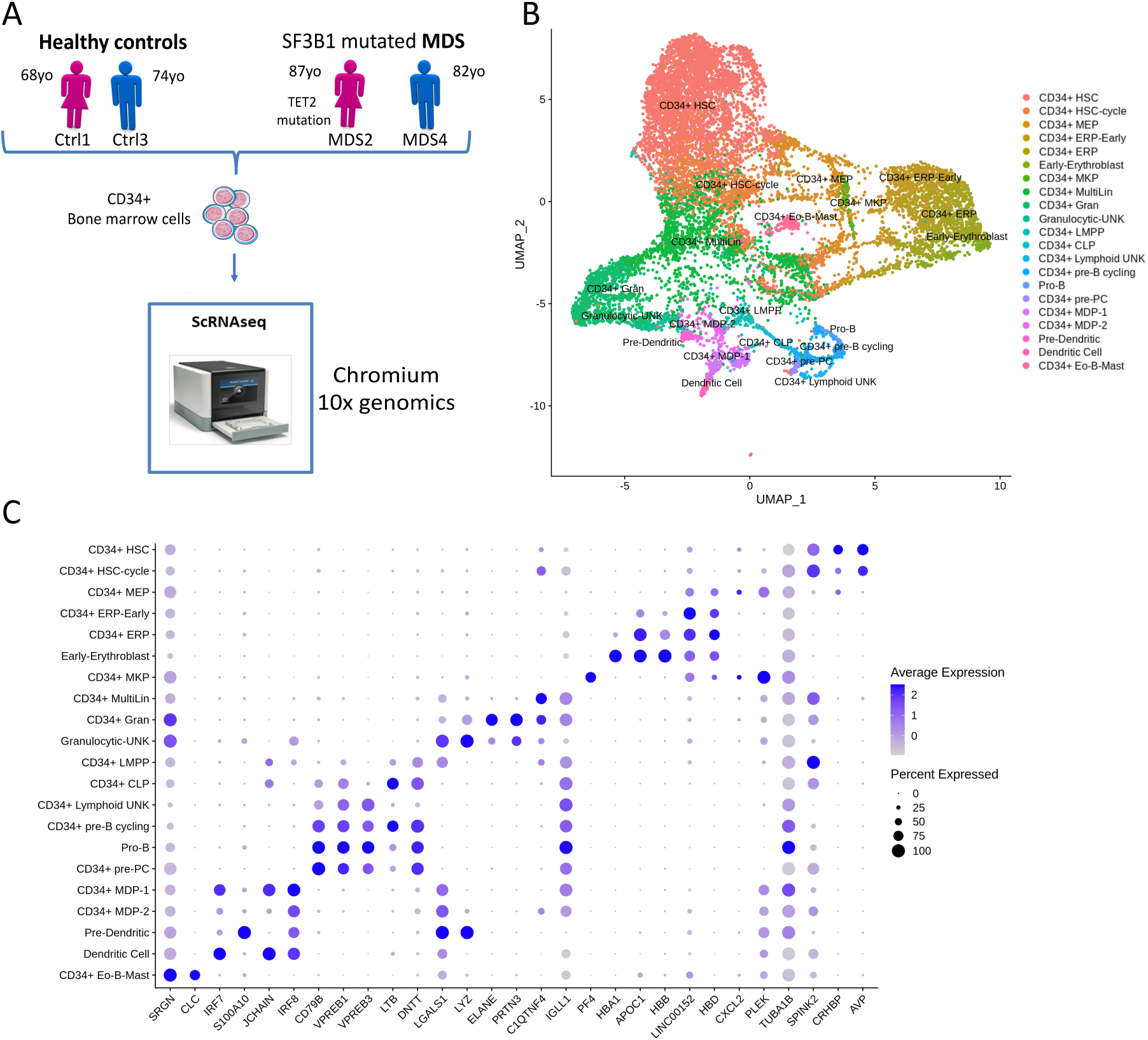
Transcriptional landscape of the HSPC compartment of SF3B1 mutated MDS and healthy elderly subjects. **A)** Outline of experimental approach. CD34+ HSPCs were isolated from bone marrow of healthy elderly controls (Ctrl1, Ctrl3) and SF3B1 mutated. scRNAseq was performed using chromium 10x genomics technology.**B-C)** Analysis of 10x Genomics scRNAseq data from 12689 cells, combining the 4 samples. **B)** UMAP of HSPC landscape. The cells annotated by SingleR are classified into 21 different subtypes, each represented by a different color. **C)** Expression values of selected marker genes for all cell sub-populations. Circle color shows mean scaled expression values and circle size represents the proportion of expressing cells per sub-populations.

**Supplementary table 2:** Clinico-biological features of patients from whom HSPCs were harvested for the scRNA-seq experiment.

| Patient | Ctrl1 | Ctrl3 | MDS2 | MDS4 |
| --- | --- | --- | --- | --- |
| Age | 68 | 74 | 87 | 82 |
| Sex | F | M | F | M |
| Diagnosis | Healthy ctrl | Healthy ctrl | MDS-RS | MDS-RS |
| R-IPSS | NA | NA | low | very low |
| Hb, g/dL | 14.3 | 15.5 | 10 | 10,6 |
| Platelets, G/L | 220 | 190 | 342 | 287 |
| Neutrophils, G/L | 1.97 | 3 | 1.4 | 3,46 |
| Monocytes, G/L | 0.47 | 0.46 | 0.1 | 0,96 |
| Lymphocytes, G/L | 1.77 | 1.63 | 0.5 | 1,74 |
| Bone marrow cellularity | NA | NA | High | Very high |
| Erythroid precursors, % | NA | NA | 60 | 52 |
| Myeloid precursors, % | NA | NA | 30 | 43 |
| Megacaryocytes | NA | NA | Presents | Numerous |
| Bone marrow blasts, % | NA | NA | 2 | 1 |
| Ring sideroblasts, % | NA | NA | 25 | 50 |
| Karyotype | NA | NA | 46,XX,del(11)(q14)[3]<br>/46,sl,?del(20)(q12)[2]<br>/46,X,del(X)(q21),add(5)(q?31),<br>del(11)(q14),add(17)(q2?2)[12]<br>/46,XX[2] | 46, XY [20] |
| Somatic mutations (VAF%) | Not detected | Not detected | TET2, p.Q769X (44%)<br>SF3B1, p.K700E (43%) | SF3B1, p.K666RK (35%) |

**Supplementary table 3:** Distribution of cellular subpopulations for each differentiation pathway in Ctrl1, Ctrl3, MDS2 and MDS4 samples.

Erythropoiesis
| Sample | CD34+<br>HSC | CD34+<br>MEP | CD34+<br>ERP-Early | CD34+<br>ERP | Early-<br>Erythroblast |
| --- | --- | --- | --- | --- | --- |
| Ctrl1 | 754 | 176 | 156 | 284 | 24 |
| Ctrl3 | 1415 | 194 | 347 | 423 | 29 |
| MDS2 | 1100 | 126 | 76 | 143 | 182 |
| MDS4 | 824 | 158 | 253 | 252 | 24 |

Granulopoiesis
| Sample | CD34+<br>HSC | CD34+<br>HSC-cycle | CD34+<br>MultiLin | CD34+<br>Gran |
| --- | --- | --- | --- | --- |
| Ctrl1 | 754 | 188 | 256 | 208 |
| Ctrl3 | 1415 | 240 | 229 | 221 |
| MDS2 | 1100 | 618 | 475 | 311 |
| MDS4 | 825 | 496 | 601 | 380 |

Dendritic differentiation
| Sample | CD34+<br>HSC | CD34+<br>HSC-cycle | CD34+<br>MultiLin | CD34+<br>LMPP | CD34+<br>MDP-2 | CD34+<br>MDP-1 | Dendritic Cell |
| --- | --- | --- | --- | --- | --- | --- | --- |
| Ctrl1 | 754 | 188 | 256 | 25 | 32 | 32 | 13 |
| Ctrl3 | 1415 | 240 | 229 | 21 | 23 | 18 | 11 |
| MDS2 | 1100 | 618 | 475 | 35 | 42 | 47 | 25 |
| MDS4 | 825 | 496 | 601 | 86 | 79 | 94 | 38 |

| Sample | CD34+<br>HSC | CD34+<br>HSC-cycle | CD34+<br>MultiLin | CD34+<br>LMPP | CD34+<br>CLP | CD34+<br>pre-B cycling | CD34+<br>pro-B |
| --- | --- | --- | --- | --- | --- | --- | --- |
| Ctrl1 | 754 | 188 | 256 | 25 | 3 | 39 | 2 |
| Ctrl3 | 1415 | 240 | 229 | 21 | 18 | 76 | 9 |
| MDS2 | 1100 | 618 | 475 | 35 | 8 | 0 | 0 |
| MDS4 | 820 | 489 | 525 | 86 | 70 | 150 | 22 |

**Figure S13:**
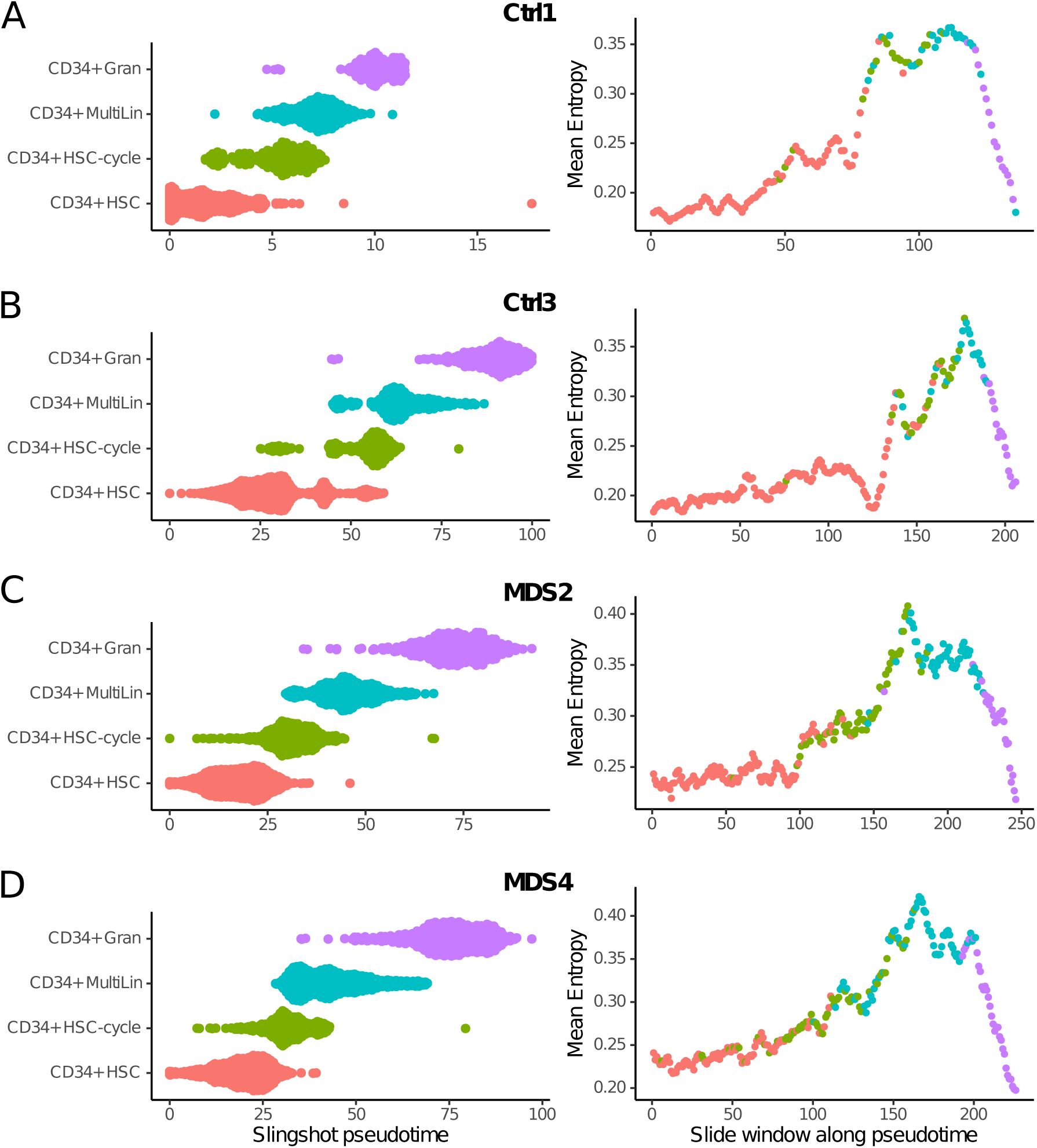
Evolution of cell-to-cell gene expression variability during granulopoiesis in elderly subjects and SF3B1-mutated MDS. For each sample individually, cell populations belonging to granulopoiesis were first selected and then ordered according to the pseudotime calculated by Slingshot. The average intercellular entropy of all genes was then calculated on a sliding window of 50 cells which moves across the pseudotime with a step of 10 cells (the color of each point on the graph correspond to the nature of the first cell in the corresponding sliding window). **A)** Ctrl1 **B)** Ctrl3 **C)** MDS2 **D)** MDS4

**Figure S14:**
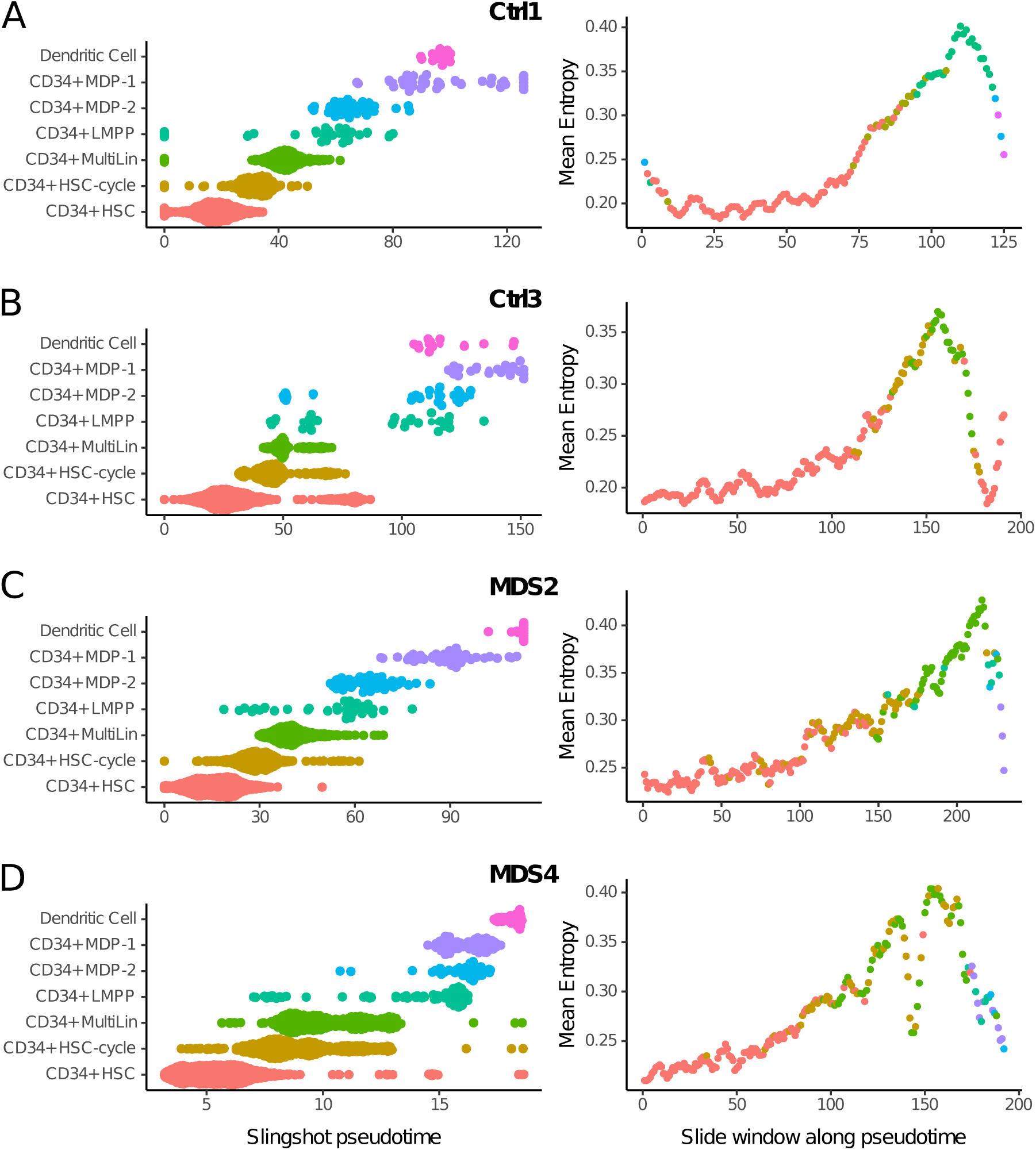
Evolution of cell-to-cell gene expression variability during dendritic differentiation in elderly subjects and SF3B1-mutated MDS. For each sample individually, cell populations belonging to dendritic differentiation were first selected and then ordered according to the pseudotime calculated by Slingshot. The average intercellular entropy of all genes was then calculated on a sliding window of 50 cells which moves across the pseudotime with a step of 10 cells (the color of each point on the graph correspond to the nature of the first cell in the corresponding sliding window). **A)** Ctrl1 **B)** Ctrl3 **C)** MDS2 **D)** MDS4

**Figure S15:**
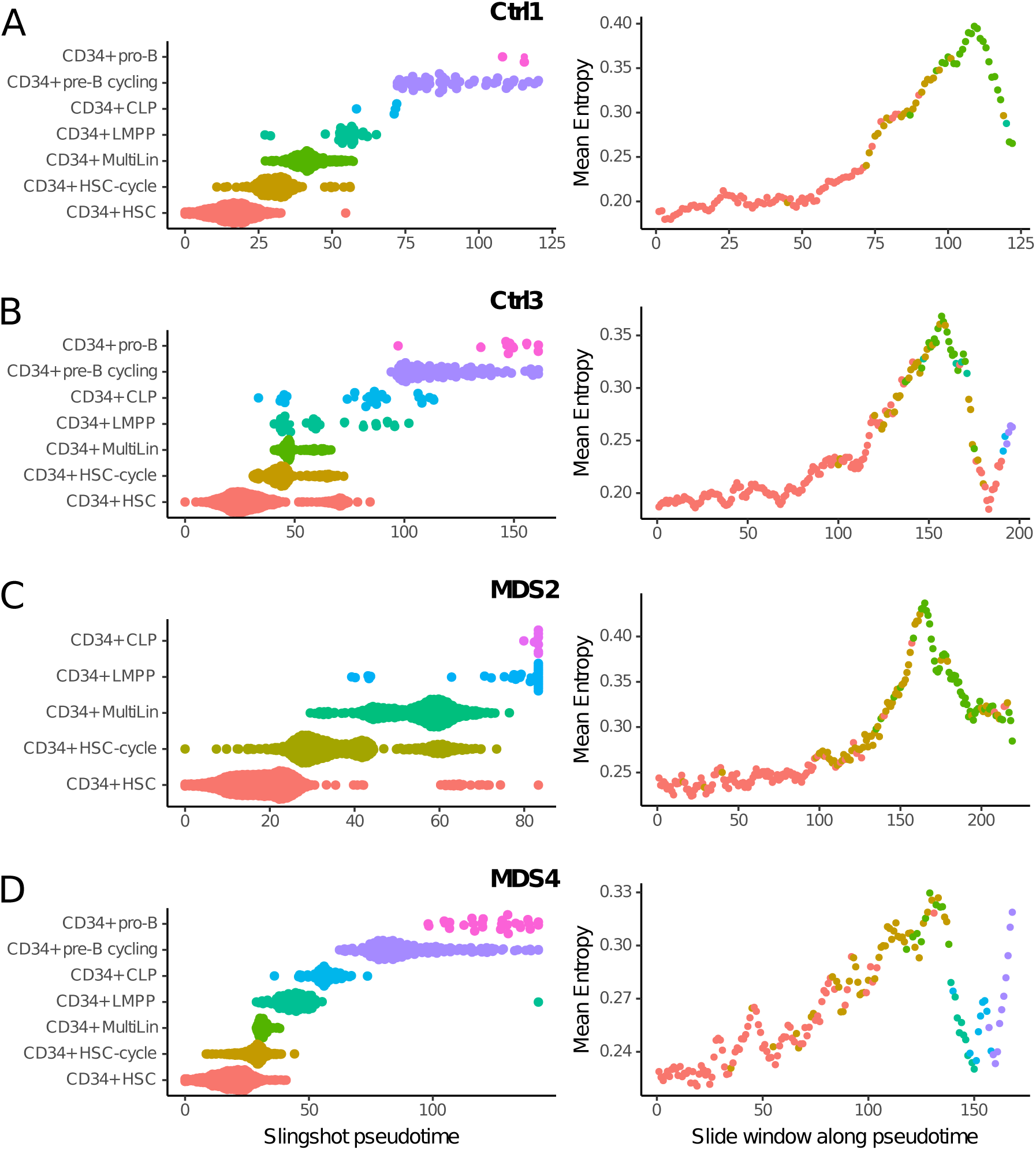
Evolution of cell-to-cell gene expression variability during B lymphopoiesis in elderly subjects and SF3B1-mutated MDS. For each sample individually, cell populations belonging to B lymphopoiesis were first selected and then ordered according to the pseudotime calculated by Slingshot. The average intercellular entropy of all genes was then calculated on a sliding window of 50 cells which moves across the pseudotime with a step of 10 cells (the color of each point on the graph correspond to the nature of the first cell in the corresponding sliding window). **A)** Ctrl1 **B)** Ctrl3 **C)** MDS2 **D)** MDS4

**Figure S16:**
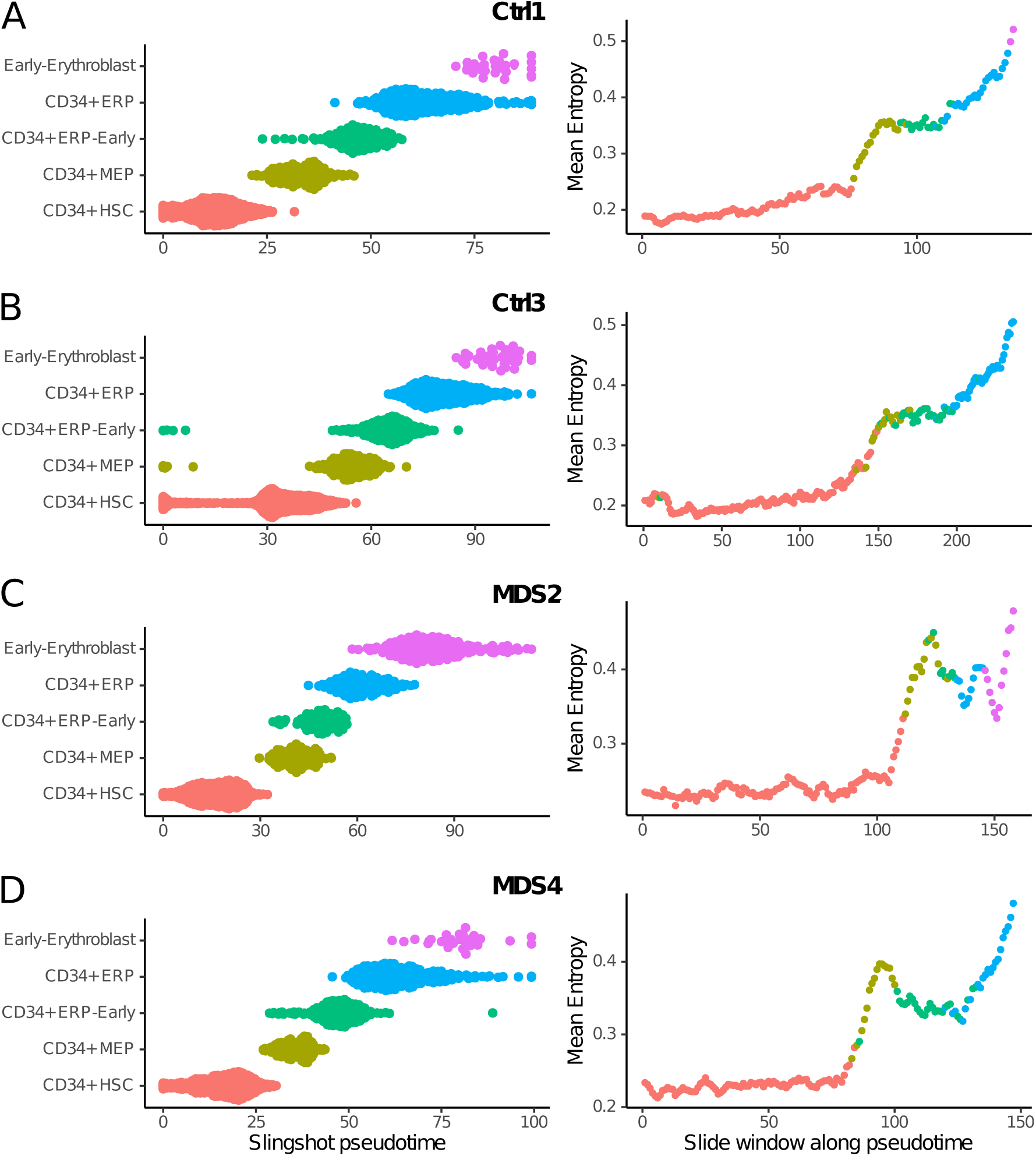
Evolution of cell-to-cell gene expression variability during Erythropoiesis in elderly subjects and SF3B1-mutated MDS. For each sample individually, cell populations belonging to erythropoiesis were first selected and then ordered according to the pseudotime calculated by Slingshot. The average intercellular entropy of all genes was then calculated on a sliding window of 50 cells which moves across the pseudotime with a step of 10 cells (the color of each point on the graph correspond to the nature of the first cell in the corresponding sliding window). **A)** Ctrl1 **B)** Ctrl3 **C)** MDS2 **D)** MDS4

